# Integrative Single-cell and Spatial Transcriptomic Analysis of Osteosarcoma Reveals Conserved and Distinct Ecosystems Across Sites and Species

**DOI:** 10.64898/2026.01.13.698472

**Authors:** Yogesh Budhathoki, Matthew V. Cannon, Troy A. McEachron, Anand G. Patel, Matthew J. Gust, Jaime F. Modiano, Dylan T. Ammons, Kathryn E. Cronise, Marissa G. Macchietto, Aaron L. Sarver, Ruben Dries, Berkley E. Gryder, Daniel P. Regan, Heather L. Gardner, Ryan D. Roberts

## Abstract

Osteosarcoma is a heterogeneous malignancy, exhibiting significant variability among patients, individual cancer cells within a tumor, and the stromal cells that compose primary and metastatic lesions. To facilitate the study of this complex disease, we compiled a unique cross-species single-cell transcriptomic dataset totaling over a million cells/nuclei from human specimens, canine specimens, patient-derived xenografts/PDX, and syngeneic mouse models at both primary (bone) and metastatic (lung) sites. Using a rigorous process for multi-species alignment and annotation, we identified six conserved tumor cell transcriptional states organized along hierarchical differentiation trajectories from progenitor to differentiated phenotypes. Parallel analysis of tumor-associated cells identified conserved macrophage, fibroblast, and endothelial populations that exhibit species– and site-specific reprogramming. Validation by mapping cell types using spatial transcriptomics revealed structured neighborhood architectures that were reproduced across multiple samples. Cell-cell interaction analysis revealed similarities and differences in tumor-host networks across primary and metastatic sites and across species. This analysis enabled pathway-specific assessment of tumor-host communication fidelity across osteosarcoma model systems relative to humans, revealing canine osteosarcoma as a more faithful model. Metastatic lung lesions, counterintuitively, exhibited more intense and complex extracellular matrix (ECM) signaling than primary bone tumors. A key example was tumor-derived fibronectin (FN1), which engages integrin and syndecan receptors on lung epithelial cells, driving a pathological mesenchymal and profibrotic state that promotes fibrotic niche formation and metastatic lung colonization. Together, this cross-species resource delineates both conserved and divergent tumor microenvironment programs, demonstrates how model-aware analyses uncover previously unrecognized tumor-host interactions, and underscores the need for therapies that co-target tumor heterogeneity and its supportive metastatic niche.

**Statement of Significance:** We show that the tumor microenvironment in human osteosarcoma patients has biological phenotypes that are conserved across both patients and species. This points towards underlying molecular mechanisms that could be therapeutically targeted.

## Introduction

Osteosarcoma is a highly heterogeneous malignancy characterized primarily by structurally complex genomes (1–6). While extensive genomic studies have documented substantial inter-and intra-tumoral diversity, fundamental questions remain about how this heterogeneity affects individual osteosarcoma cell states, tumor cell lineage plasticity, and environmental tumor-host interactions that drive disease progression and metastasis, a major determinant of patient mortality. These challenges are compounded by the widespread use of experimental models whose ability to faithfully recapitulate the cellular hierarchies and tumor–host interactions of human osteosarcoma has not been systematically evaluated (7–10).

Single-cell transcriptomic technologies are uniquely well-positioned to address these questions by directly measuring cell states, lineage relationships, and microenvironmental interactions at high resolution. Several studies have applied single cell RNA sequencing (scRNA-seq) to osteosarcoma, establishing important precedents for characterizing tumor heterogeneity (2,7–10). However, these efforts have been limited by small cohort sizes that used study-specific nomenclature, single-institution sampling lacking the numbers needed to reach rigorous conclusions, limited understanding of similarities between models and human disease, and the absence of spatial context. Moreover, accurately identifying malignant cells and harmonizing tumor cell states across studies and model systems remain significant unresolved challenges.

To overcome these limitations, we assembled a multi-institutional, multi-species osteosarcoma dataset comprising 775,441 quality-controlled single-cell/nucleus transcriptomes from 129 distinct samples representing both primary and metastatic sites, spanning human patients, canine patients, patient-derived xenografts (PDX), and syngeneic/genetically engineered mouse models. Together with the validation datasets and normal-tissue comparators, the resource encompasses over a million single cells/nuclei, representing unprecedented scope and scale for the study of a specific rare disease. The scale and multi-site, multi-species nature of this dataset was deliberately designed to facilitate the identification of tumor and tumor-associated host cell subsets with unprecedented detail and rigor. Leveraging the salient characteristics of this unique dataset, we developed a unified framework for the robust and reproducible identification of tumor cells, harmonized annotations to enable confident identification of orthologous populations, and conducted systematic comparisons across species, anatomical contexts, and experimental platforms. We then extended these findings using spatial transcriptomic profiling to provide structural context for tumor and tumor-associated cell populations.

Using this integrative single-cell and spatial framework, we delineate conserved and divergent osteosarcoma tumor programs, define the composition and organization of the tumor microenvironment, and quantitatively assess the fidelity with which commonly used model systems reproduce specific and essential elements of the human disease. Surprisingly, metastatic lung lesions exhibited markedly more pronounced and more complex extracellular matrix-derived signaling compared with primary bone tumors. Within these interactions, we identify tumor-derived fibronectin (FN1) as a key mediator of reprogramming within the metastatic niche, engaging syndecan and integrin receptors on lung epithelial cells to induce inflammatory, mesenchymal, profibrotic programs reminiscent of pulmonary fibrosis (11–13). Spatial transcriptomic analyses reveal conserved intra-lesional architectures that support distinct tumor cell states across samples.

Together, this work establishes a cross-species, spatially-resolved framework for studying osteosarcoma heterogeneity, model fidelity, and tumor-microenvironment interactions, providing a foundational model for defining heterogeneous subpopulations and a resource for discovery, accurate biological inference, and scientifically rigorous translational targeting.

## Materials and Methods

### Data Acquisition

The data used in this study were acquired from Colorado State University (14) (CSU), Gene Expression Omnibus (7) (GSE152048, GSE162454, GSE217792, GSE283885), National Cancer Institute (15) (NCI), Nationwide Children’s Hospital (NCH), St. Jude Children’s Research Hospital (SJ), University of Minnesota (UoM), and Tufts University (16) (TU). All of the mouse samples came from treatment-naive animals, whereas human patient samples consisted of a combination of patients who underwent standard care. Samples used in this study underwent either single-cell, single-nucleus RNA sequencing or Visium spatial (15) transcriptomics at the specific institution from which they were sourced. Only lung samples were used for the metastatic group, and the primary group consisted of a combination of different sites, including tibia, femur, and humerus. All of the sample details can be found in the sample detail table (Table 1).

### Sample Preparation

Sample preparation details, including tissue dissociation, single-cell and single-nucleus RNA library preparation, and sequencing for each sample/dataset, can be found below and the respective publication each dataset was sourced from.

#### National Cancer Institute Datasets

All solutions were prepared under sterile conditions and kept on ice before use. For each sample, four 15 mL tubes, one gentleMACS C tube, and two DNA LoBind microcentrifuge tubes were pre-labeled and chilled on ice, along with additional tubes for sample weighing and counting. One 70 µm and one 30 µm cell strainer were allocated per sample. NP-40 lysis buffer was prepared in advance (10 mM Tris-HCl, pH 7.5, 10 mM NaCl, 3 mM MgCl₂, 0.1% NP-40, 1 mM DTT, 1 U/µL RNase inhibitor), and 1.9 mL was dispensed into each C tube and kept on ice.

Frozen tissue samples were processed depending on the sample type. Small fragments were weighed using pre-chilled microfuge tubes on dry ice, chopped into fine pieces with sterile razor blades on a dry-ice–chilled 10 cm dish, and transferred directly into the C tube containing NP-40 lysis buffer. Large frozen tissues were first fractured by striking the foil-wrapped sample with a pestle in a liquid-nitrogen–chilled mortar, weighed on dry ice, and minced prior to transfer. For bone or calcified tissues, pulverization in liquid nitrogen was performed until the material reached a fine powder, which was then transferred into a C tube. OCT-embedded samples were first trimmed to remove excess embedding compound, rinsed briefly in cold PBS to dissolve remaining OCT, minced into small pieces, and immediately transferred into NP-40 lysis buffer without weighing to minimize thawing.

Samples were homogenized using a gentleMACS Octo Dissociator (Miltenyi Biotec) with the “4C_nuclei_1” program (∼5 min). Homogenates were briefly centrifuged to collect contents and incubated on ice for 5 min. The lysate was filtered through a 70 µm MACS SmartStrainer into a pre-chilled 15 mL tube, followed by a rinse of the C tube with 950 µL NP-40 lysis buffer to recover residual material. Samples were centrifuged at 500 × g for 5 min at 4°C. The supernatant was transferred to a new 15 mL tube, leaving ∼100 µL above the pellet, and stored at −80°C for optional downstream proteomics.

The nuclei pellet was gently overlaid with 900 µL PBS containing 1% BSA and RNase inhibitor (0.2–1 U/µL), incubated on ice for 5 min, resuspended, and centrifuged again at 500 × g for 5 min at 4°C. The wash was repeated once more, passing the nuclei through a 30 µm MACS SmartStrainer into a fresh 15 mL tube. Nuclei were counted on a Countess II FL using AO/PI staining, with appropriate dilutions (1:10 or 1:20) in PBS + 1% BSA + RNase inhibitor as needed. The concentration of viable (GFP⁺) nuclei was used to calculate the input volume required to obtain at least 30 µL containing ∼5 × 10³ nuclei/µL.

Calculated nuclei volumes were centrifuged at 500 × g for 5 min at 4°C in swinging-bucket rotors to ensure efficient pelleting. The supernatant was discarded, and nuclei were resuspended in 100 µL of 0.1× lysis buffer (10 mM Tris-HCl, 10 mM NaCl, 3 mM MgCl₂, 0.1% NP-40, 0.1% Tween-20, 0.01% digitonin, 1% BSA, 1 mM DTT, 1 U/µL RNase inhibitor) and incubated on ice for 2 min. Following incubation, 950 µL wash buffer (10 mM Tris-HCl, 10 mM NaCl, 3 mM MgCl₂, 1% BSA, 0.1% Tween-20, 1 mM DTT, 1 U/µL RNase inhibitor) was added, and samples were centrifuged at 500 × g for 5 min at 4°C. The resulting nuclei pellet was gently resuspended in Diluted Nuclei Buffer (1× Nuclei Buffer supplemented with 1 mM DTT and 1 U/µL RNase inhibitor) to achieve a final concentration of 5 × 10³ nuclei/µL and transferred to low-bind microfuge tubes kept on ice.

Nuclei suspensions were immediately submitted for 10x Genomics Chromium Single-Nucleus Multiome (RNA + ATAC) library preparation. Remaining nuclei were imaged using an EVOS microscope at 10× magnification to confirm integrity and morphology, then centrifuged at 500 × g for 5 min at 4°C and frozen at −80°C for archival. All reagents and buffers were freshly prepared using molecular-grade water and sterile filtered before use.

snMultiome capture and library prep: The quality and the concentration of the single nuclei suspension were first checked by staining the nuclei with AO/PI fluorescent dye (F23001, Logos Biosystems) and using the Luna-FL Dual Fluorescent Cell Counter (Logos Biosystems). The quality was determined by checking the viability (less than 5% viability), clumping of nuclei (clumping is a sign of compromised nuclei), and average size. The samples with more than 5% viable cells were usually not captured, but given the nature of the osteosarcoma samples with bone fragments, a few samples had a higher false-positive background. In those cases, the loading concentrations were calculated using the number of dead cells. The nuclei suspension was either diluted or concentrated to get the nuclei concentration in the range of 4000-5000 nuclei/µl. The quality and the concentrations were checked again, and then the single nuclei capture with a target recovery of 7000 nuclei was performed by following the 10X Genomics User Guide (CG000338). Briefly, the open chromatin regions of the DNA were first transposed, and the adapter sequences were added to capture the DNA fragments for the ATAC library. The transposed nuclei, along with RT mix and beads, were added onto the assembled NextGEM Chip J and loaded immediately to the Chromium Controller for GEM generation and barcoding. After the Post-GEM cleanup and pre-amplification PCR, 10X barcoded DNA fragments and the 10X barcoded full-length cDNA from mRNA in the pre-amplified product were used to prepare the ATAC library and the GEX library, respectively.

Sequencing: The concentration and average size of the ATAC and GEX final libraries were quality checked by Agilent 2100 Bioanalyzer (Agilent). Equimolar concentrations of the libraries from all the samples were pooled and sequenced using the Illumina Sequencer, NovaSeq-6000. The ATAC and GEX libraries were pooled separately and sequenced at the sequencing depth of 50,000 reads per nucleus. Sequencing Parameters: GEX Library – 28|10|10|90, R1|I1|I2|R2 and ATAC library – 50|8|24|49, R1|I1|I2|R2)

Data Processing: Raw BCL files were first demultiplexed and then aligned to the 10x Genomics provided human reference genome (refdata-cellranger-arc-GRCh38-2020-A-2.0.0). The demultiplexing and alignment were done using the cellranger-arc pipeline, version 2.0.0. The count matrix files generated for each sample were further used for secondary data analysis.

#### Nationwide Children’s Hospital

Lungs harvested were processed using the human tumor dissociation kit (Miltenyi Biotec, 130-095-929) with a gentleMACS Octo Dissociator with Heaters (Miltenyi Biotec, 130-096-427) for 3′ single-cell analysis. For single-nucleus analysis, cryopreserved samples were isolated and permeabilized for single-cell gene expression sequencing following the Nuclei Isolation from Complex Tissues for Single Cell Multiome ATAC + Gene Expression Sequencing protocol (10x Genomics, CG000375 Rev C). Following dissociation into single cells or nuclei, 3′ RNA-seq libraries were prepared using Chromium Single Cell 3′ RNA-seq system (10x Genomics) with the Reagent Kit v3.1 (10x Genomics, PN-1000121) according to the manufacturer’s instructions. cDNA library quality and quantity were determined using an Agilent 2100 bioanalyzer using the High Sensitivity DNA Reagents kit (Agilent, 5607-4626) and then sequenced on an Illumina NovaSeq 6000.

#### University of Minnesota

Experimental subjects: dogs with treatment-naive appendicular osteosarcoma and no evidence of metastasis were recruited into the study with owner consent (UMN IACUC protocol 2201-39751A). An incentive of $500 was available to defray costs of histopathology for eligible dogs.

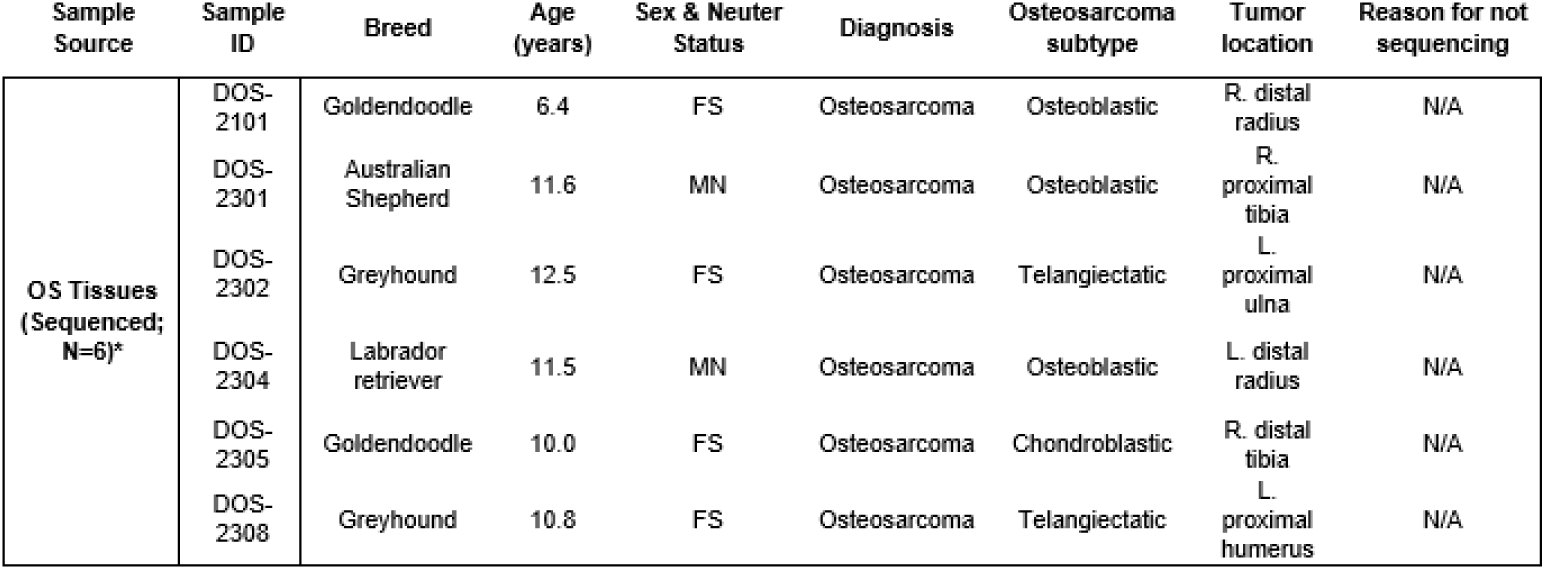

| Sample Source | Sample ID | Breed | Age (years) | Sex & Neuter Status | Diagnosis | Osteosarcoma subtype | Tumor location | Reason for not sequencing |
| --- | --- | --- | --- | --- | --- | --- | --- | --- |
| OS Tissues (Sequenced; N=6)* | DOS-2101 | Goldendoodle | 6.4 | FS | Osteosarcoma | Osteoblastic | R. distal radius | N/A |
|  | DOS-2301 | Australian Shepherd | 11.6 | MN | Osteosarcoma | Osteoblastic | R. proximal tibia | N/A |
|  | DOS-2302 | Greyhound | 12.5 | FS | Osteosarcoma | Telangiectatic | L. proximal ulna | N/A |
|  | DOS-2304 | Labrador retriever | 11.5 | MN | Osteosarcoma | Osteoblastic | L. distal radius | N/A |
|  | DOS-2305 | Goldendoodle | 10.0 | FS | Osteosarcoma | Chondroblastic | R. distal tibia | N/A |
|  | DOS-2308 | Greyhound | 10.8 | FS | Osteosarcoma | Telangiectatic | L. proximal humerus | N/A |

Sample processing: The affected, amputated limb was immediately transferred to the histopathology processing laboratory of the University of Minnesota Veterinary Diagnostic Laboratory for processing. The musculature and connective tissue were dissected away from bone and gross images were captured on a faxitron (see Makielski, Mol Ther Oncolytics 2023 for reference). The bone was sagittally sectioned with a bone saw. A section of the tumor material that came in contact with the saw was dissected away and discarded, and tumor sections were recovered and saved as independent blocks for further processing. A single block was immersed in transport media (DMEM supplemented with 10% FBS and 0.1% Primocin) for single cell recovery.

The tumor section was disrupted by repeated mechanical dissociation using a sterile scalpel, scissors, and forceps in a 10 cm Petri dish containing sterile phosphate buffered saline. Single cells were separated from stromal elements by sequential filtration through sterile 100 µm, 40 µm and 30 µm mesh filters. In samples with excess blood contamination, red blood cells were eliminated through a 1-step lysis protocol.

Cell number and viability were initially determined based on uptake of trypan blue using a Countess II cell counter and confirmed based on propidium iodide uptake and integration using a Luna cell counter. Cells were suspended at the concentration recommended by 10x Genomics for single cell capture in the Chromium platform. Cell captures, library creation, and sequencing were done at the University of Minnesota Genomics Center using standard methods credentialed for compatibility with the 10X platform. Data delivery required passing standard QC metrics for all samples.

#### Tufts University

All procedures were performed under sterile conditions to prevent contamination. Prior to cell processing, a bead bath was set to 37°C, and all subsequent washes were conducted at room temperature. RPMI medium supplemented with 10% FBS (RPMI C10) was pre-warmed for cell thawing. Wash buffer (1× DPBS containing 0.04% BSA; 0.08 g BSA per 200 mL) was sterile-filtered and stored at 4°C. Pre-sorting buffer (1× DPBS with 10% FBS) was prepared and chilled on ice, while pre-coating buffer (1× DPBS with 20% FBS) and collection buffer (PBS with 20% FBS or as specified) were freshly prepared before sorting.

Cryopreserved samples were rapidly thawed in a 37°C bead bath for 2–3 minutes and immediately transferred to a 50 mL conical tube using wide-bore pipette tips. Each cryovial was rinsed with 1 mL of warm RPMI C10, and the rinse was combined with the cell suspension. Warm RPMI C10 (20 mL) was slowly added, and the suspension was gently mixed by inversion. Cells were centrifuged at 150 × g for 10 minutes at room temperature, and the supernatant was carefully removed, leaving approximately 200 µL of residual liquid. Based on the initial cell concentration and viability, cells were resuspended in RPMI C10 to achieve a final concentration of approximately 2.5 × 10⁶ cells/mL. Osteosarcoma samples were passed twice through a 70 µm strainer, whereas DLBLC samples were filtered twice through a 40 µm strainer, followed by centrifugation under identical conditions. The pellet was resuspended in RPMI C10 to the same target concentration. Cell concentration and viability were assessed using acridine orange/propidium iodide (AO/PI) staining on a Cellometer K2. Samples with a viability exceeding 85% proceeded directly to downstream processing, while those below this threshold were subjected to fluorescence-activated cell sorting (FACS) to remove nonviable cells.

For washing, viable cells were centrifuged at 300 × g for 5–10 minutes, the supernatant was discarded, and the pellet was gently resuspended in 1 mL of DPBS containing 0.04% BSA using wide-bore pipette tips. This wash step was repeated twice. The final pellet was resuspended in DPBS + 0.04% BSA at approximately 2.5 × 10³ cells/µL, and viability was reassessed using AO/PI staining. The suspension was diluted as needed to a final target concentration of 700–1200 cells/µL in DPBS + 0.04% BSA. The cell suspension was maintained on ice and immediately subjected to 10x Genomics single-cell library preparation protocols.

For samples requiring flow sorting, cells were centrifuged at 150 × g for 10 minutes at room temperature and resuspended in chilled pre-sorting buffer at a ratio of 100 µL per 1 × 10⁶ cells. 7-AAD (5 µL per 100 µL sample; 0.25 µg) was added to each sample, followed by a 10-minute incubation in the dark. Sorting collection tubes were pre-coated with PBS containing 20% FBS, after which the coating solution was removed and replaced with collection buffer. Cells were sorted using purity mode under low pressure (100 µm nozzle), with collection tubes maintained on ice. Post-sorting, cells were centrifuged at 150 × g for 10 minutes at 4°C, the supernatant was discarded, and the pellet was resuspended in DPBS + 0.04% BSA. Cell concentration and viability were confirmed by AO/PI staining, and the suspension was adjusted to a final concentration of 700–1200 cells/µL. Sorted cells were kept on ice and immediately processed for 10x Genomics single-cell library construction.

Detailed methods can be found in: https://www.biorxiv.org/content/10.64898/2026.01.05.696411v1.full

#### Colorado State University

OS pulmonary metastases were obtained by a veterinary pathologist within 1-4 hours post-euthanasia from dogs submitted for autopsy (Colorado State University) or from canine OS patients undergoing metastasectomy (C. Thomson, Ethos Veterinary Health) with informed owner consent and stored in complete DMEM on ice until processing. Samples were washed with PBS, minced into 3-5 mm pieces, and then suspended in collagenase II (Gibco #17101-015) at 250U/mL in HBSS with calcium and magnesium (Gibco #24020-117). Samples were dissociated in a 37°C water bath for a total of 45 minutes while vortexing at maximum speed every 5 minutes. Cell suspensions were then passed through a 70 μm cell strainer, washed with PBS, and centrifuged at 400 x*g*. The pelleted cells were resuspended in 2-5 mL of Ammonium-Chloride-Potassium lysis buffer, incubated at room temperature for 3 minutes, diluted in 12 mL of PBS, centrifuged at 400 x*g*, washed with PBS, and then centrifuged again at 400 x*g*. Pelleted cells were then resuspended in 2-5 mL of 0.04% UltraPure BSA (Thermo Fisher Scientific #AM2616) in PBS, passed through a 70 μm cell strainer, counted on a hemacytometer, and readjusted to a concentration of 700-1,200 cells/μL. Samples with a viability of at least 70% were captured on a Chromium iX (10x Genomics) with a target of 5,000 cells per sample. Library preparation was performed using the Chromium Next GEM Single Cell 3’ v3.1 Dual Index Kit (10x Genomics) according to the manufacturer’s protocol. Libraries were sequenced by Novogene Corporation using a NovaSeq 6000 (Illumina) with a target depth of 50,000 150bp paired-end reads. Raw FASTQ sequencing reads were aligned using the CellRanger (10x Genomics) pipeline to the canine UU_Cfam_GSD_1.0 genome assembly (Ensembl release 112) appended with the ROS_Cfam_1.0 Y chromosome and Canfam 3.1 mitochondrial genome. For the primary tumor samples, methods can be found at https://www.nature.com/articles/s42003-024-06182-w.

#### St. Jude Children’s Research Hospital

Patient samples accrual: Flash-frozen tissue samples were obtained through the St. Jude Children’s Research Hospital Biorepository after approval by the St. Jude Children’s Research Hospital Institutional Review Board (protocol ID XPD17-183).

Xenograft sample accrual: Flash-frozen fragments of osteosarcoma orthotopic patient-derived xenografts (PDXs) were obtained via request through the St. Jude Childhood Solid Tumor Network (PMID# 26068307; https://cstn.stjude.cloud/).

Tissue processing: Flash-frozen tumor fragments were processed for single-nucleus RNA-sequencing (snRNA-seq) according to the TST extraction protocol (Slyper et al., 2020; PMID# 32405060). Flash-frozen tumor fragments (approximately 50-100 mg) were incubated for 10 minutes in 1 mL TST buffer (73 mM sodium chloride, 5 mM Tris [pH 8.0], 0.5 mM calcium chloride, 10.5 mM magnesium chloride, 0.01% bovine serum albumin, 0.03% Tween-20) while mincing with Noyes spring scissors. Nuclei suspensions were then filtered through a 40 mm strainer, followed by rinsing of the strainer with an additional 1 mL of TST buffer. Nuclei suspensions were diluted with 3 mL of ST buffer (73 mM sodium chloride, 5 mM Tris [pH 8.0], 0.5 mM calcium chloride, 10.5 mM magnesium chloride) and centrifuged for 5 min at 500 x*g* at 4 degrees C. The nuclei pellet was then resuspended in 100-500 mL ST buffer and filtered through a 40 mm strainer prior to snRNA-seq. A step-by-step protocol is available at: https://dx.doi.org/10.17504/protocols.io.bhbcj2iw

Osteosarcoma sample processing for single-cell/nucleus RNA-sequencing: scRNA-seq and snRNA-seq were performed using version 3 of the 10x Genomics Single Cell 3’ RNA Expression Solution kits. Ten thousand nuclei were input into the 10x Chromium controller for droplet partitioning with barcoded beads. Barcoded libraries were generated according to manufacturer instructions. Each library underwent paired-end sequencing (30,000 paired end reads/cell) on an Illumina NovaSeq X Plus sequencer and processed using bclfastq to generate: Read 1 – 26 nucleotides, Read 2 – 100 nucleotides, Index – 8 nucleotides.

Data Availability: St. Jude osteosarcoma snRNA-seq data will be available through GEO accession code GSE293805.

### scRNA-seq Data Analysis Pipeline

We used Cell Ranger (v7.2.0, 10x Genomics) to align the reads to a joint human/mouse (GRCh38/mm10) genomic reference and generate gene count data per cell. For dogs, sequencing reads were aligned using the Cell Ranger to the canine UU_Cfam_GSD_1.0 genome assembly (Ensembl release 112), appended with the ROS_Cfam_1.0 Y chromosome and Canfam 3.1 mitochondrial genome. Where needed for cross-species analysis, dog genes were converted into human orthologs using Ensembl’s BioMart database (17). We analyzed the data in R using the Seurat (18) package. We used strict quality control measures for each sample to eliminate low-quality cells and potential doublets. We applied sample-specific cutoffs for maximum unique molecular identifiers (UMIs) per cell to remove doublets. Any cell with fewer than 1000 UMIs and any sample with fewer than 100 cells were filtered out of the analysis to maintain quality. We filtered out dead and dying cells using a maximum value cutoff for the percentage of mitochondrial reads per cell. We then randomly down-sampled cells to have a maximum of 5000 cells per sample, merged the datasets for each group, and used Seurat to normalize and scale the data for downstream analysis. 2000 variable genes were used for dimension reduction with principal component analysis, followed by Uniform Manifold Approximation and Projection (UMAP) embedding. Unsupervised clustering was achieved using “FindNeighbors” and “FindClusters” functions in Seurat using the top 30 PCA dimensions. Clustering resolution was determined using the Clustree (19) R package, and the Harmony (20) package was used for the integration and visualization of data.

### Tumor cell identification

We used five different methods to identify tumor cells: SCEVAN (21), scATOMIC (22), CopyKAT (23), ScanBit (24), and truth-set reference-based cell type annotation. Ultimately, cancer cells from humans, mice, and dogs were identified using the truth-set reference-based cell type annotation method that leverages the SingleR (25) package. Fleiss’ kappa was calculated to measure agreement among different groups (GitHub code).

### Stromal cell type annotation

The stromal cell types were initially annotated using a combination of species and site-specific references within the celldex package and references derived from publicly available resources utilizing SingleR. Stroma for primary human (and dog) datasets was initially annotated using a combination of HumanPrimaryCellAtlasData, and MonacoImmuneData within the celldex R package using SingleR (25). Stroma for metastatic human datasets was initially annotated utilizing a combination of HumanPrimaryCellAtlasData, MonacoImmuneData, and Human Lung Cell Atlas (HCLA)(26). The stroma for the primary mouse and xenograft datasets was initially annotated using the MouseRNAseqData and ImmGenData from the CellDex package, utilizing SingleR (25). The stroma for the metastatic mouse and xenograft datasets was annotated utilizing a combination of curated mouse lung cell atlas, MouseRNAseqData, and ImmGenData using SingleR. Using the AUCell(27) package, we also looked at the expression of cell type markers outlined in several publications (28,29) and PanglaoDB (30) to assess cell type assignment accuracy. Because the stromal cells in our datasets were tumor-associated, we used these initial SingleR annotations to subdivide these stromal cells into four broader groups, namely 1) epithelial and endothelial, 2) immune myeloid, 3) immune lymphoid, and 4) mesenchymal. We then reprocessed each group as explained above. After reclustering, each group was subjected to AUCell analysis using cell type-specific markers, and each cluster was manually assigned a final cell type annotation. We used differentially expressed genes identified using the DESeq2 (31) package (with batch variables included as covariates) and known cell type-specific markers to aid the final cell type annotations. The cell types were annotated with increasing granularity into four levels: level 0, level 1, level 2, and level 3. Level 3 consists of the cell type annotation with the highest granularity, and level 0 is the annotation with the lowest.

### Tumor sub-population annotation

We optimized clustering resolution for each dataset using the Clustree (19) algorithm, selecting the resolution at which cluster branching stabilized. To ensure the process underlying the Harmony integration did not overfit and create artifactual clusters, we reprocessed the data from each sample separately, then performed a correlation analysis to compare clusters identified within individual samples to those defined in the integrated dataset. Clusters identified when samples were processed individually showed strong correlations with those in the integrated dataset, validating the tumor subpopulations identified.

We identified differentially expressed genes in the tumor cells using the DESeq2 (31) package (with batch variables as covariates). We then performed gene set enrichment analysis (GSEA) using the fgsea package (32) on the Reactome pathways from the MSigDB (32) package to identify altered pathways for each cluster. After optimization of clustering resolution using the Clustree package, tumor clusters representing at least 5% of the total tumor cells and showing distinct pathway enrichment were classified as independent subpopulations; smaller clusters were merged with the most similar larger group. We performed RNA velocity using scVelo(33) to visualize differentiation trajectories as outlined below. We also performed regulon analysis using the SCENIC (27) package to infer and identify underlying regulatory enhancers to aid our cancer subpopulation annotation. Default parameters were used to run sample-by-sample regulon analysis. P values were computed using a Wilcoxon rank-sum test and adjusted for multiple testing using the Benjamini–Hochberg procedure.

The annotated Seurat objects containing the human dataset were used as a reference to annotate both primary and metastatic osteosarcoma scRNA-seq datasets from the Alex Lemonade Foundation repository (34).

### RNA Velocity Analysis

RNA velocity analysis was conducted to model the evolution of subpopulations within tumor cells and tumor-associated macrophages. Loom files containing each sample’s gene expression and splicing information for every cell were generated with velocyto (35) v0.17.17, using each sample’s BAM file and cell barcodes that passed previous quality control as input. The loom files were used to create anndata (v0.10.7) objects for analysis in Python v3.12.9. We added metadata from Seurat objects, including cell type labels, Seurat clusters, and UMAP, PCA, and FDL embeddings. To comply with scVelo’s (33) modeling assumptions, only single-cell samples were included, except for xenograft samples, where only single-nucleus data were available and thus used for velocity analysis. RNA velocity modeling and analysis were performed using Scanpy (36) v1.11.0 and scVelo v0.3.4 on both tumor and myeloid populations. Velocity was modeled using “scv.tl.velocity,” which produced velocity vectors for each gene and cell. These velocity vectors were then transformed into a cell-by-cell matrix of transition probabilities using “scv.tl.velocity_graph.” This transition matrix was projected into UMAP space using “scv.tl.velocity_embedding_stream.” Cell type trajectory inference was performed with Partition-based Graph Abstraction (PAGA) using “scv.tl.paga,” which computes connectivity between cell types and leverages RNA velocity directionality to quantify cell state transitions.

### Doublet Detection

Doublet detection was undertaken as a quality control step using the Demuxafy (37) suite, which incorporates DoubletDetection, scDblFinder, and SCDS. To assess Demuxafy’s accuracy, a subset of known doublets was established by identifying droplets from PDX samples that contained at least 200 total RNA reads, with a mixture of human and mouse reads (50±10% mapped to the mouse genome and 50±10% to the human genome).

Further assessment of Demuxafy was carried out by merging stromal cells from human patients with human tumor cells from PDX models into a single Seurat object. The resulting data was written to disk using the write10xCounts function from the DropletUtils(38) R package and was provided as input for Demuxafy. Since no new doublets are expected in this merged dataset, the proportion of false-positive doublet calls allows estimation of Demuxafy’s precision.

We also sought to understand if the Interactive tumor cells were doublets by applying the Demuxafy suite (39), which combines DoubletDetection (40), scDblFinder (41), and SCDS (42). Unsurprisingly, scDblFinder and SCDS reported portions of the Interactive cells as putative doublets, whereas DoubletDetection was less consistent. The overall result from the doublet detection was unclear and inconsistent across each dataset. To test the reliability of these results, we utilized a known doublet truth set, comprising PDX cells with both mouse and human genes (cross-species doublets). Strikingly, Demuxafy identified only ∼10% of these as doublets, suggesting that doublet calls are unreliable in cases of mixed transcriptional profiles.

### Spatial Analysis

Eight preprocessed 10x Visium spatial transcriptomics datasets from the Eigenbrood et. Al (15) study were downloaded as RDS files and analyzed with Seurat in R. Since the files were already preprocessed, no additional quality control was necessary. To estimate the cell type composition of spatial spots in the Visium data, we used the RCTD method via the SpaceXR (43) R package, using patient metastatic sequencing data and cell-type annotations from this study as a reference. To assess the spatial correlation of cell types, we utilized Lee’s (44) L statistic implemented in the spdep R package.

### CellChat Analysis

Mouse genes were converted to their human orthologs using the NicheNet convert_mouse_to_human_symbols function within the nichenetr (45) package. Dog genes were converted into human orthologs using Ensembl’s BioMart (17) database, producing one harmonized gene space for downstream ligand–receptor analysis. For each dataset, we constructed a sample-by-sample CellChat (46) object using cell type identities. To create this sample-by-sample cellchat object, we first subsetted the Seurat object to a maximum of 100 cells per cell type at annotation level 3. The human CellChat ligand–receptor database CellChatDB.human was used to infer communication inference. Before communication inference, we applied subsetData to retain expressed signaling genes and ran identifyOverExpressedGenes and identifyOverExpressedInteractions to select candidate ligand and receptor pairs. Communication probabilities were computed using computeCommunProb and aggregated at the signaling pathway level with computeCommunProbPathway and aggregateNet. netAnalysis_computeCentrality is used in CellChat to quantify the relative importance or influence of each cell group (cell type or cluster) within the inferred cell–cell communication network. Default parameters were used while applying each function within the CellChat package.

To identify dominant site-specific ligand–receptor signaling pathways, we first computed average communication probabilities across all samples for each site. Only pathways that were present in humans were retained across all species. Pathways were then stratified by site (primary vs. metastatic). We filtered for pathways whose sum in all species was less than the human average in each (primary and metastatic) site, and ranked them by their total summed interaction probability across all species. The top 40 pathways (or fewer than 40, if fewer pathways passed the filtering) per site were retained for downstream visualization.

To enable cross-species comparison, data must ultimately be converted to a human format; however, this translation introduces inherent limitations due to differences in immune signaling, ligand–receptor usage, and antigen-presentation machinery. For example, MHC-II gene divergence complicates direct equivalence between mouse and human systems.

For cross-species fidelity analyses, we correlated the cellchat results of humans (gold standard) to each model (dog, mouse, PDX) in each primary and metastatic site. Correlations were computed using the receiver cell type, ligand, and receptor information from the CellChat result. We calculated pairwise Spearman correlations between humans and each of the corresponding model systems. Functional signaling modules were defined by using pathways enriched in each site. Modules were annotated based on the dominant biological themes of their constituent pathways, yielding seven major categories: ECM Remodeling, Cell Adhesion, Angiogenesis, Immune Modulation, Stemness, Neuronal, and Metabolic.

## Data Availability

All code used in this analysis is available on GitHub at https://github.com/kidcancerlab/Osteo_atlas. The raw data used in the creation of this dataset are publicly available through the following repositories:

– GEO accession numbers GSE152048, GSE162454, GSE217792, GSE283885.
– The ALSF Single-cell Pediatric Cancer Atlas Portal project numbers SCPCP000017, SCPCP00018, and SCPCP00023.
– Human lung cell atlas data from EGA numbers EGAD00001006126, EGAD00001006127, and EGAD00001006128.
– Healthy mouse lung reference data from ENA number PRJNA632939.
– dbGaP accession numbers phs004431.v1.p1 and phs003569.v1.p1.

Processed and annotated R-loadable datasets are hosted on FigShare at doi:10.6084/m9.figshare.31029559.

## Results

To systematically chart conserved and divergent cellular programs in osteosarcoma and to benchmark the fidelity of commonly used model systems, we assembled a comprehensive cross-species scRNA-seq compendium from human, canine, PDXs, and syngeneic mouse models (Fig. 1A). The resulting dataset unified the analysis of primary and metastatic lesions across two distinct disease sites and three different species. To capture the full spectrum of osteosarcoma across clinical presentations, species, methods (whole-cell and single-nucleus), and model systems, we consolidated data from six major research centers and supplemented it with publicly available datasets. Samples were obtained from Nationwide Children’s Hospital, the National Cancer Institute, St. Jude Children’s Research Hospital, Colorado State University, the University of Minnesota, and Tufts University, as well as from public repositories (GSE152048, GSE162454, GSE217792, GSE283885) (Fig. 1A). A total of eight datasets were constructed based on species/experimental model and site of disease: human primary, human metastatic, dog primary, dog metastatic, PDX primary, PDX metastatic, mouse primary, and mouse metastatic. Each dataset was processed uniformly but independently, using a Seurat v5-based pipeline (47) (Fig. 1B), which included alignment to common reference transcriptomes and parameter-optimized Harmony (20) integration (Supplementary Figs. 1).

**Figure 1.**
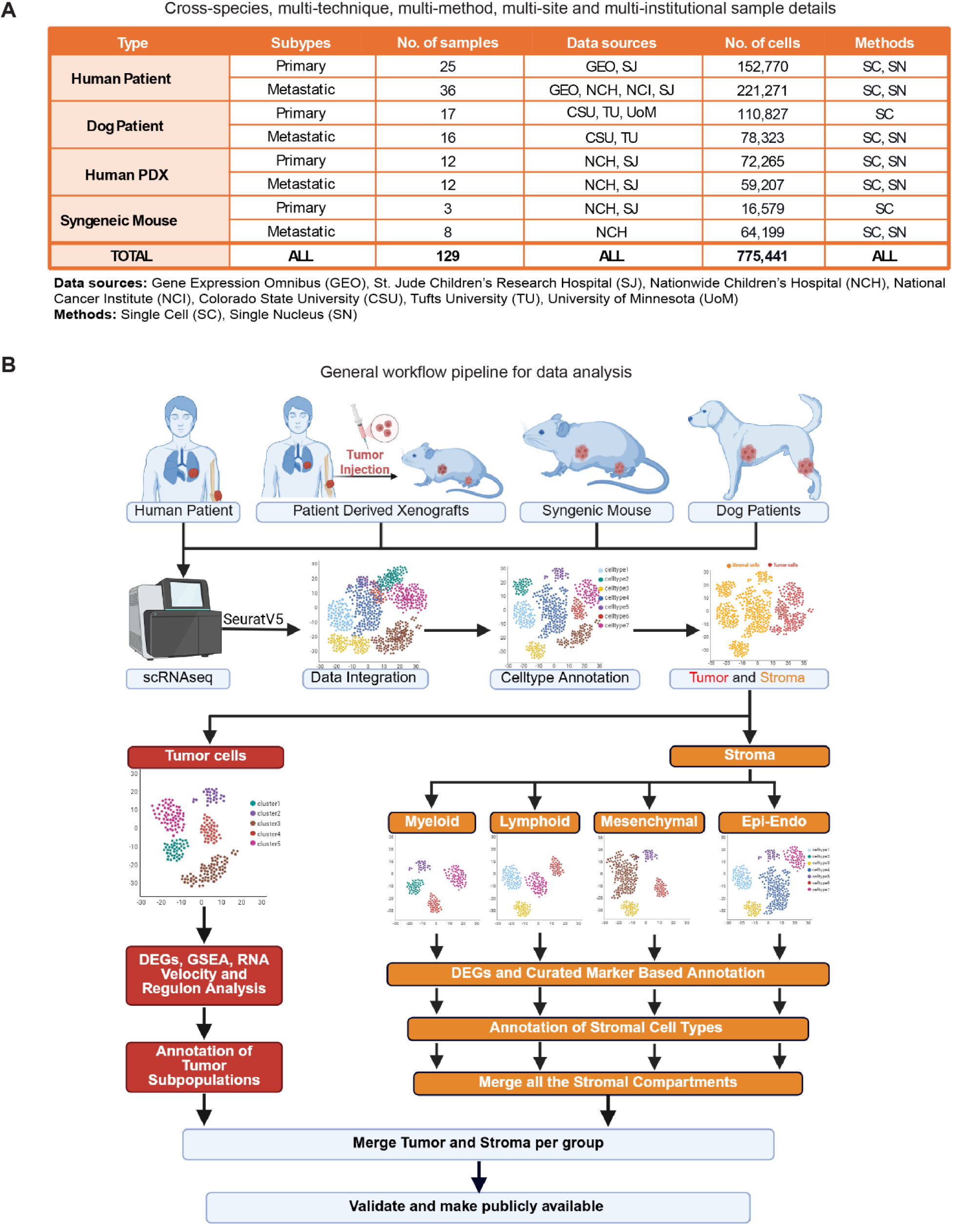
Cross-species sample overview and analytical workflow. A) Summary table detailing sample characteristics across datasets, including sample type, anatomical site, total number of samples, source, total number of cells, and sequencing methodology. B) Schematic of the overall analytical workflow outlining data integration, tumor and stromal cell identification, and cell type annotation steps used to construct the cross-species osteosarcoma atlas.

### Novel pipelines leveraging cross-species truth sets and SNV-based evolutionary algorithms effectively resolve tumor-stromal ambiguity

To maximize integration accuracy and minimize confounding effects arising from analyses that combine highly heterogeneous malignant cells with stromal cells (which exhibit relatively low inter-subject heterogeneity), we first aimed to accurately isolate the malignant cells from each sample. We applied three widely used methods, SCEVAN (21), scATOMIC (22), and CopyKAT (23), to our human and mouse datasets. We found striking variability and discordance in the malignant cell calling by these algorithms (Fig. 2A and Supplementary Fig. 2). Notably, a quantitative evaluation of inter-rater reliability using Fleiss’ kappa indicated that agreement between the methods was only modestly better than would be expected by chance (Fig. 2A). This inconsistency was particularly concerning in the metastatic datasets.

**Figure 2.**
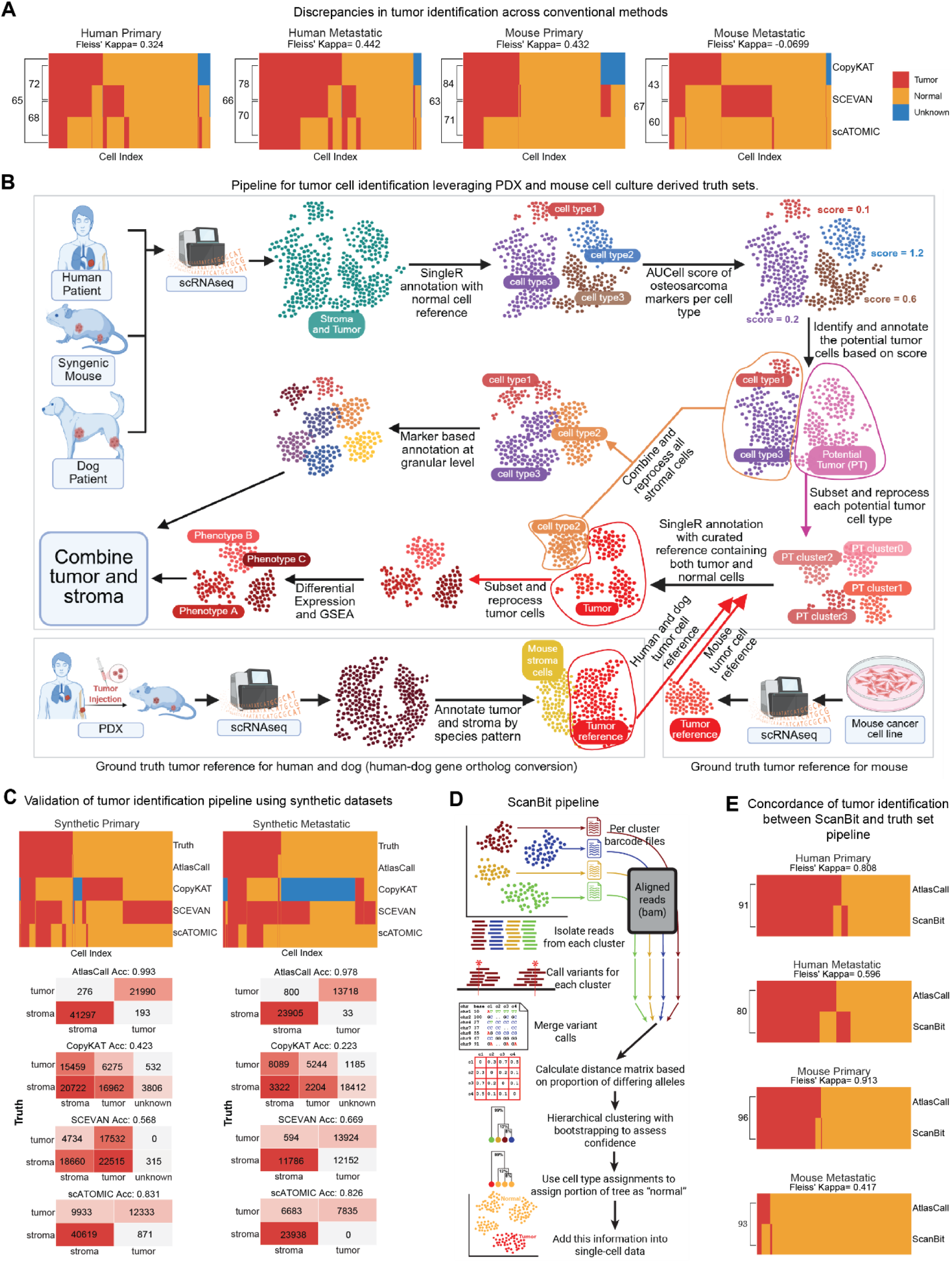
Cross-species truth set and SNV-based algorithm effectively resolve tumor–stromal ambiguity. A) Heatmap depicting discrepancies in tumor identification across conventional methods (CopyKAT, SCEVAN, scATOMIC), with corresponding Fleiss’ Kappa coefficients for each dataset (human primary, human metastatic, mouse primary, and mouse metastatic). B) Overview of the pipeline developed for accurate tumor cell identification, integrating osteosarcoma-specific markers and leveraging patient-derived xenograft (PDX) and mouse culture–derived truth set references in combination with relevant species and tissue-specific stromal reference data. C) Schematic of the ScanBit pipeline, an orthogonal validation method used to confirm tumor cell classification accuracy. D) Heatmap comparing tumor and normal cell classifications obtained using the truth set–based pipeline, CopyKAT, SCEVAN, and scATOMIC against ground-truth annotations, demonstrating improved accuracy of the truth set-based pipeline across both primary and metastatic samples. E) Heatmap showing concordance between tumor–stromal classifications obtained from the orthogonal ScanBit pipeline and the truth set–based approach, validating robust and consistent tumor identification.

To address this, we sought to establish a disease-relevant process that could reliably distinguish malignant cells from stromal components. We quickly realized that the structural characteristics of our dataset facilitated the identification of some tumor cells with very high confidence, which might be used to create disease-specific malignant cell truth sets. For instance, in the PDX datasets, human malignant cells and mouse stromal cells could be cleanly segregated by species-specific transcript alignments. These tumor cells could be isolated and combined with non-malignant disease datasets to create a robust human malignant truth set (Fig. 2B). Similarly, scRNA-seq profiles from cell lines derived from several of the murine models could facilitate the creation of a mouse malignant truth set (Fig. 2B). These truth sets formed the basis of a two-step classification framework in which initial assignments, generated using healthy stromal references and area under the curve (AUC) scores (48) for specific markers of osteosarcoma, were refined using these species– and site-matched malignant/stromal truth set references (Fig. 2B).

To evaluate the precision of this new pipeline, we created benchmark tests using semi-synthetic samples in which the identities of malignant and stromal cells were already known (Fig. 2C). We randomly selected four PDX samples, two from primary sites and two from metastatic sites, each containing at least 5,000 cells. Importantly, these samples were not part of the reference truth set used for malignant cell identification. To mimic the fibrotic and reprogrammed tumor microenvironment observed in osteosarcoma, we combined primary PDX malignant cell data with normal bone stroma data from a human fracture nonunion dataset (49) (GSE217792) and metastatic PDX malignant cell data with normal lung stroma data from an idiopathic pulmonary fibrosis dataset (50) (GSE283885).

Applying our new method to these semi-synthetic datasets, where the true identities of malignant and stromal cells were established, we achieved at least 95% accuracy in identifying malignant cells (Fig. 2C), significantly outperforming SCEVAN (21), scATOMIC (22), and CopyKAT (23) (Fig. 2D and Supplementary Fig. 2A). For real human and mouse datasets, malignant and stromal compartments were separated with high accuracy (Supplementary Fig. 2). Interestingly, when processing canine samples, a simple dog-to-human ortholog mapping followed by annotation using the human malignant truth set reference produced excellent results, obviating the need to create a canine-specific truth set (Supplementary Figs. 2B and 2C).

To validate malignant cell identification using an orthogonal method, we also applied ScanBit (24) (Figs. 2D, 2E), a new single-nucleotide variation (SNV)-based approach for malignant cell identification, to these same samples. Identities produced by the ScanBit algorithm and the truth set pipeline showed a very high degree of inter-rater consistency (Fig. 2E), providing convincing evidence for the accuracy of our malignant cell identification. This approach was robust and reliable across human, canine, and mouse datasets in both primary and metastatic samples. This new method has broad applicability: it can be used to identify tumor cells in any malignancy for which a truth set can be confidently created.

### Six conserved tumor subpopulations define osteosarcoma across most species and disease sites

After confidently separating malignant and non-malignant cells, we retained 410,791 confidently annotated malignant cells across datasets. Our next objective was to characterize the transcriptionally-defined subpopulations within the malignant cell compartment. To do this, we first pooled all tumor cells from each species (human, canine, and mouse), then optimized and validated integration and clustering parameters for each tumor cell dataset to ensure that every cluster represented a transcriptionally distinct entity (Supplementary Fig. 3).

Next, we aimed to annotate these tumor subpopulations using Differential Gene Expression (DEG) analysis and Gene Set Enrichment Analysis (GSEA). Because Harmony addresses batch effects at the clustering and visualization level but does not correct expression values, we used a pseudobulk approach to aggregate counts at the sample level for each cluster, then perform differential expression analysis with DESeq2 (31). We then performed GSEA (51) to identify biologically meaningful patterns within these differentially-expressed genes using MSigDB (32) Reactome pathways as a reference. This identified six biologically distinct tumor subpopulations, which we designated as Proliferative, chondroid-osteoid matrix-associated (COMA), Fibrogenic, Basal progenitor, Metabolically Primed (MP) progenitor, and Interactive (Fig. 3 and Supplementary Fig. 4). Importantly, we detected these subpopulations across nearly all datasets, and identified shifts in proportion between primary and metastatic sites (Supplementary Fig. 5). Of note, the COMA population was absent in genetically engineered mouse models (Fig. 3), for reasons that became clear in downstream analyses. Proliferative cells showed cell-cycle activation (MKI67, TOP2A); COMA cells expressed hypoxia and angiogenesis markers (BNIP3, VEGFA); Fibrogenic cells were enriched for extracellular matrix genes (POSTN, TNC); Progenitor states exhibited metabolic or stem-like programs (RPL genes; CD36, PDGFD); and Interactive cells expressed immune lineage markers (CSF1R, PTPRC) (Fig. 3).

**Figure 3.**
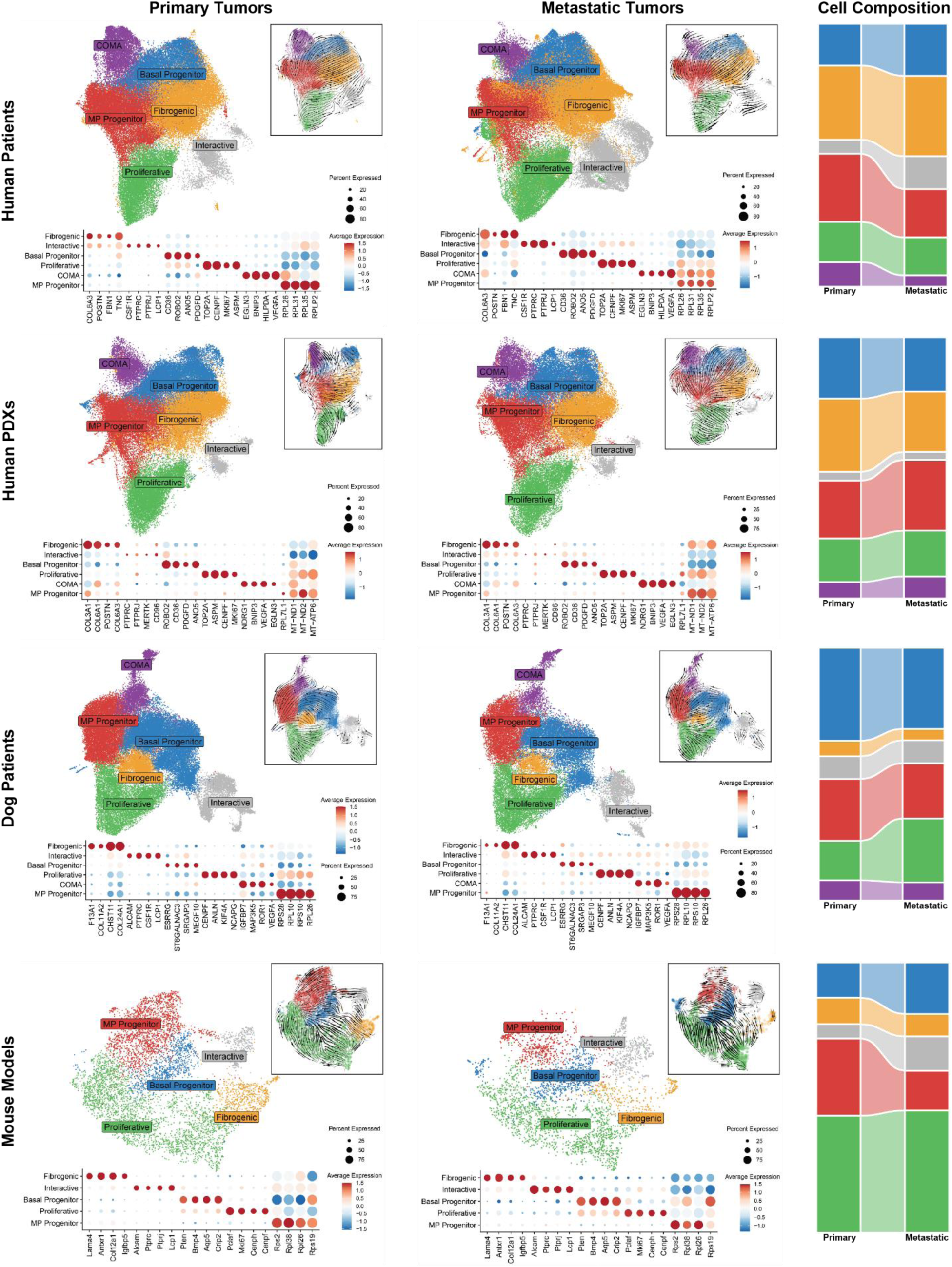
Six conserved tumor subpopulations define osteosarcoma across species and disease sites. UMAP embeddings (DimPlots) depict the distribution of tumor subpopulations identified in human, human PDX, dogs, and key syngeneic mouse models datasets at primary and metastatic sites. Corresponding dot plots display the expression of key marker genes defining each tumor subpopulation. Insets within each panel show RNA velocity projections illustrating dynamic transcriptional trajectories and potential lineage relationships among tumor states. The Sankey plot summarizes changes in the proportional distribution of tumor subpopulations between primary and metastatic sites across human, PDX, dog, and mouse datasets, highlighting conserved and site-specific tumor state transitions.

Surprisingly, several of the genes that distinguished the Interactive cluster from the others are typically restricted to cells having macrophage or lymphoid lineages (along with pro-inflammatory genes and other cytokines). These features of the Interactive subpopulation raised concern for potential stromal contamination and/or tumor-stromal doublets. However, the presence of the Interactive population within the patient-derived xenograft (PDX) tumor compartment, where tumor identity was known with certainty (defined by expression of human gene sequences), provided confidence that this cluster is, in fact, *bona fide* malignant cells (Fig. 3). Furthermore, the Interactive cells clustered distinctly from TAMs, indicating a distinct transcriptional phenotype (Supplementary Fig. 6). Marker expression reinforced this distinction: Interactive cells expressed tumor markers at levels consistent with other tumor populations, while also expressing subsets of typically immune-restricted genes at lower levels (Supplementary Figs. 6C-6D). We also verified that interactive cells were not doublets (Supplementary Fig. 7). Having found that the Interactive tumor cells were likely true tumor cells and not stromal contamination or doublets, we next sought to understand the developmental relationships of these tumor cell states.

### Tumor cell subpopulations exhibit developmental relationships that are conserved across models and species

To investigate the developmental relationships of these distinct tumor cell states, we first employed RNA velocity analysis (33). The resulting trajectories unsurprisingly showed origination primarily within the MP progenitor, Basal progenitor, and Proliferative subpopulations, with subsequent progression toward more differentiated cell types, including the Fibrogenic, COMA, and Interactive clusters (Fig. 3). These results reinforced the appropriateness of our annotations, highlighting a hierarchical organization of tumor cell phenotypes connected by dynamic transitions that may underlie tumor evolution and adaptation.

We next performed gene regulatory analysis using SCENIC (48), to define tumor subpopulation-specific transcriptional programs agnostic of disease site. This analysis revealed distinct regulon activity across tumor cell states. Relative to other tumor cells, fibrogenic cells showed strong activation of stress-response and inflammatory regulons, including ATF3, CEBPB, CEBPD, CREB5, FOS/FOSB, and STAT1(52–55), along with developmental regulators such as MEIS1 (56), consistent with a matrix-associated phenotype (Supplementary Fig. 8A). In contrast, Proliferative cells were characterized by enrichment of cell cycle–associated regulons, including E2F1, E2F7, and PBX3 (57–59) alongside repression of inflammatory regulators such as CEBPB and SPI1 (Supplementary Fig. 8A). The Interactive subpopulation exhibited pronounced activation of immune-associated regulons, including SPI1, ETS1, IKZF1, and STAT3 (54,60–62), indicating engagement with immune signaling pathways. Basal progenitor cells showed activation of early developmental transcriptional regulators, including PBX1 and HNF4G (63,64), coupled with suppression of immune and proliferation-related programs including SPI1, IKZF1, and E2F7 (Supplementary Fig. 8A). The MP progenitor subpopulation displayed a progenitor-like regulatory profile marked by activation of MAZ and YY1 (65,66) and repression of differentiation-associated factors such as ZEB1 (67) (Supplementary Fig. 8A). COMA cells showed partial overlap with Basal progenitor, including repression of E2F1 and IKZF1, but exhibited no significantly enriched regulons overall. Together, these results demonstrate that osteosarcoma tumor subpopulations are defined by distinct and stable transcriptional regulatory programs that align with their inferred lineage hierarchy.

We then examined how regulon activity changes between primary and metastatic sites within each tumor subpopulation. This analysis revealed a subpopulation-specific enrichment for conserved and distinct transcriptional regulation across tumor sites. The metastatic tumors were characterized by broad suppression of stress-response and AP1–associated regulons (active in primary tumors), including JUN, JUNB, JUND, FOS, ATF4, and KLF2 (52,53,68) as well as inflammatory regulators such as SPI1, IRF9, and STAT1 (54,55,60) (Supplementary Fig. 8B). In contrast, metastatic tumors showed consistent activation (repressed in primary tumors) of developmental and chromatin-associated programs, including ETV6, PBX1, NFIA, TCF7L2, and HMGA2 (69–71), alongside EMT– and plasticity-associated regulators such as ZEB1, ETS1, and TEAD1 (72,73) (Supplementary Fig. 8B). These changes were not uniform across states but showed clear subpopulation specificity, with the strongest activation of plasticity-associated programs observed in progenitors, Interactive, and Proliferative compartments, while Fibrogenic and COMA states exhibited more context-dependent modulation.

### Stromal cell subpopulations are largely conserved across species but differ from primary to metastatic sites

Pivoting to address the stromal cell compartment, we took a similar approach to that used for the tumor cells, subsetting and processing the 341,162 stromal cells into distinct species– and site-defined datasets, then cross-referencing to ensure consistent treatment and annotation (Supplementary Figs. 9-10). To facilitate accurate and detailed annotation, we first divided cells into myeloid, lymphoid, endothelial–epithelial, and mesenchymal compartments and treated each subset independently.

Myeloid heterogeneity was a prominent feature, with single-cell RNA sequencing resolving dendritic cells, mast cells, monocytes, neutrophils, and distinct tumor-associated macrophages (TAMs) subtypes that were conserved across sites and species. Aside from minor species-specific differences in markers, at the broad level, the major cell types were alveolar macrophages, dendritic cells (DC1, DC2, and plasmacytoid DCs) (28,29), and TAMs (Fig. 4). TAMs displayed functional specialization: fibrogenic TAMs, IFN TAMs, inflammatory TAMs, scar-associated TAMs, proliferating TAMs, and osteoclast-like TAMs (Fig. 4). Additional cell types included mast cells and monocytes (Fig. 4). The clear presence of osteoclast-like TAMs in lung metastases was notable given their characteristic restriction to bone tissues. RNA velocity (33) analysis revealed that TAMs differentiation trajectories predominantly originated from less differentiated, general TAMs and progressed toward specialized states such as scar-associated, inflammatory, IFN, and osteoclast TAMs (Fig. 4).

**Figure 4.**
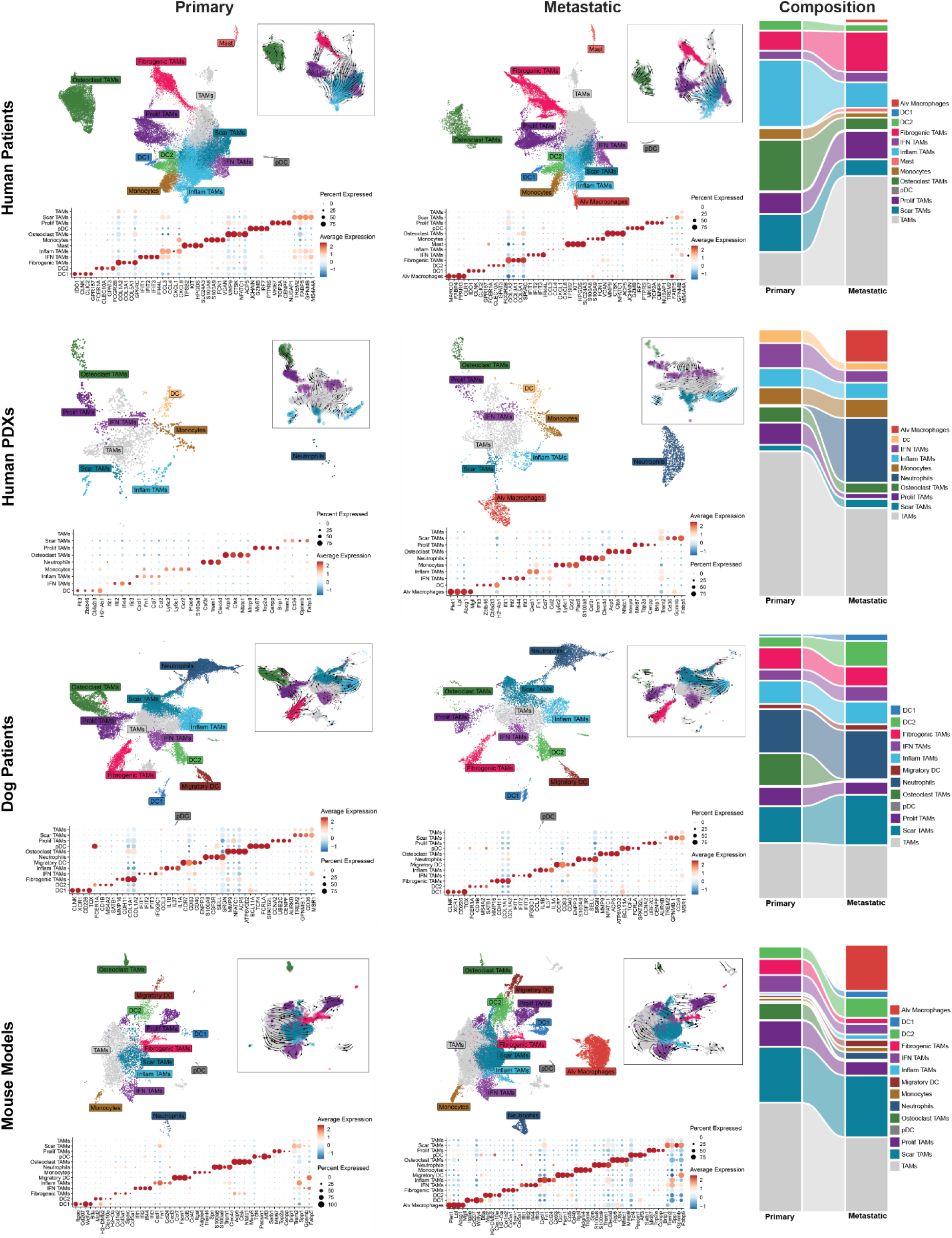
Myeloid cell subpopulations are conserved across species but shift between primary and metastatic sites. UMAP embeddings (DimPlots) illustrate the distribution of myeloid cell types, including tumor-associated macrophage (TAM) subpopulations, across human, human PDX, dogs, and syngeneic mouse datasets at primary and metastatic sites. Corresponding dot plots display the expression of key marker genes defining each myeloid subpopulation. Sankey plots summarize changes in the proportional composition of specific myeloid cell types and TAM subtypes between primary and metastatic sites across datasets, highlighting conserved myeloid programs and site-specific remodeling of the immune microenvironment.

The proportions of these TAM subsets shifted depending on disease site: primary sites favored inflammatory and osteoclast-like TAMs, while metastatic niches, especially in the lung, were enriched for scar-associated and immunosuppressive phenotypes. Syngeneic mouse and canine models best reflected the diversity of human TAMs; PDX models preserved core TAM identities but lacked fibrogenic TAMs, Mast cells, and dendritic subtypes, which may be due to differences in inter-species tumor-host interactions and constraints imposed by immunodeficiency.

We identified a variety of lymphoid cell states representing the gamut of T and B cell function, such as Naïve, CD4+ and CD8+ T cells, memory and plasma B cells, NK cells, proliferating T cells, and regulatory T cells (Supplementary Fig. 10A). Syngeneic mouse and dog stroma matched the granularity of human compartments. In contrast, PDX models exhibited markedly reduced lymphoid diversity, dominated by innate lymphoid and NK-like cells, reflecting the absence of an intact immune microenvironment and underscoring the limitations of PDXs for studying tumor–immune interactions (Supplementary Fig. 10A).

Endothelial and epithelial compartments were structurally conserved across organisms. AT1 and AT2 cells, capillary aerocytes, ciliated cells, and multiple endothelial subtypes, such as activated, arterial, capillary, lymphatic, proliferating, and venous, were consistently detected across datasets (Supplementary Fig. 10B). Canine and murine tumors preserved similar architecture, with mice showing unique subsets such as club cells, erythroblasts, and transitional alveolar cells. PDX models consolidated endothelial subtypes and showed attenuated endothelial diversity (Supplementary Fig. 10B).

The mesenchymal compartment included fibroblast subtypes such as adipogenic CAFs (adiCAFs), antigen-presenting CAFs (apCAFs), matrix (mCAFs), myofibroblastic CAFs (myCAFs), and proliferating CAFs (pCAFs), alongside pericytes, mesenchymal stem–like cells (MSCs), osteoblasts, chondrocytes, neuronal-like, and smooth muscle-like cells (Supplementary Fig. 10C). The architecture paralleled other solid tumor atlases but revealed species-dependent molecular divergence. Specific subsets exhibiting species-specific enrichment included neuronal-like mesenchymal cells in humans and mice, muscle-like mesenchymal cells in PDX, and chondrocytes or stressed osteoblasts, which were confined to human tumors. Others have reported some of these cell types previously. Specifically, neuronal features have been implicated in disease aggressiveness (74–77). Some of these subsets are conserved across human, mouse, and PDX tumors, providing direct evidence of neural, muscle, and basal-like host cell reprogramming within the osteosarcoma microenvironment.

### Reference-based annotation of spatial datasets reveals clear patterns of organization of tumor and stromal subpopulations

To assess whether the cellular subpopulations identified in our single-cell analysis is reliable and exhibited higher-order spatial organization within metastatic lung lesions, we used our rigorously annotated datasets as reference in a re-analysis of the Alex Lemonade Foundation repository single cell (34) and spatial transcriptomics data recently published by Eigenbrood et al (15) (Fig. 5 and Supplementary Fig. 11-12). Spatial deconvolution of cell-types with SpaceXR (43) revealed a highly organized and biologically meaningful spatial architecture. The deconvoluted maps showed a clear segregation between tumor and stromal compartments, with sharp spatial boundaries that were consistent across biological replicates (Fig. 5 and Supplementary Fig. 12).

**Figure 5.**
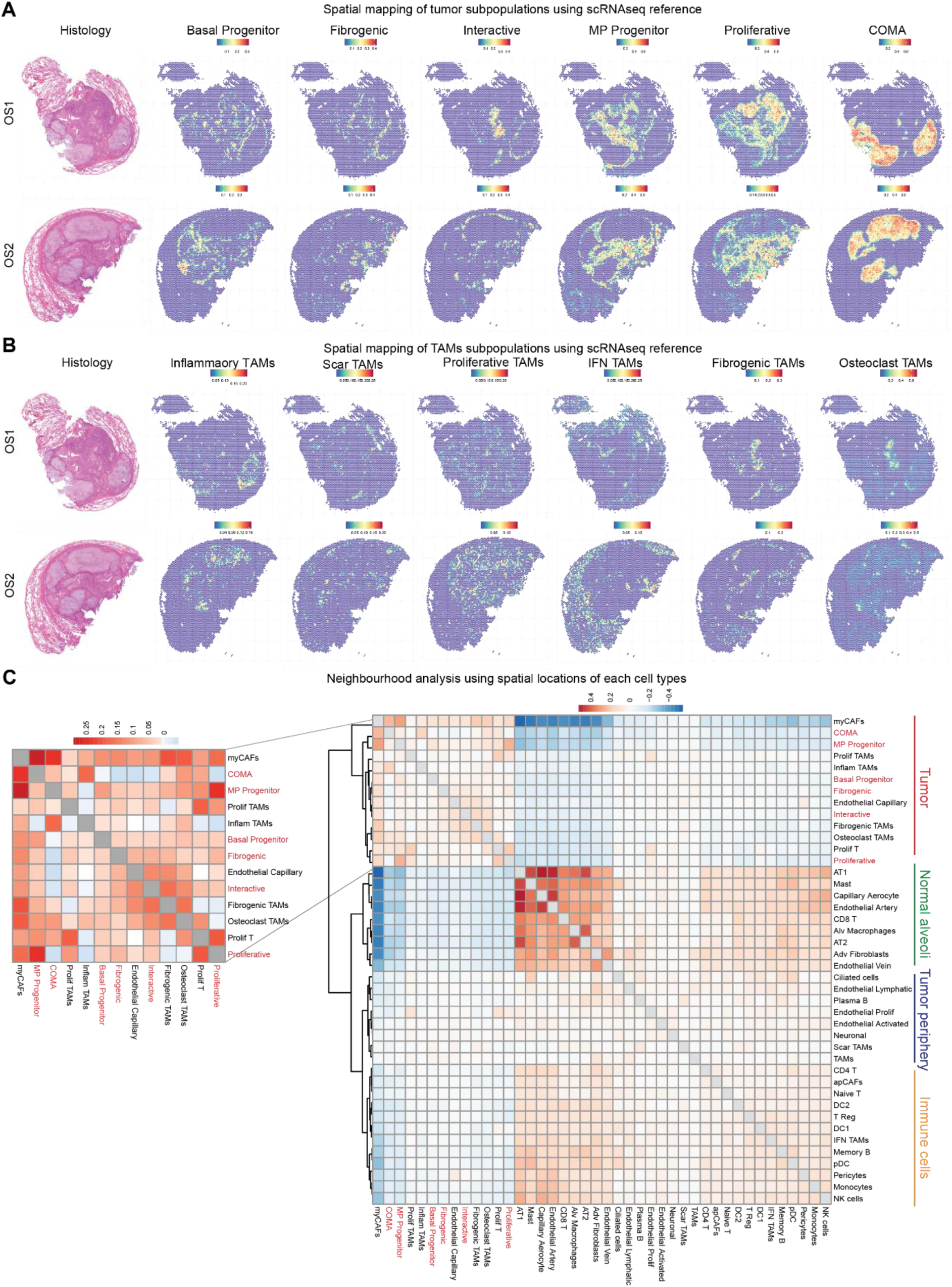
Atlas-based annotation of spatial datasets reveals organized and conserved patterns of tumor and stromal architecture. A) Spatial mapping of tumor subpopulations using the scRNA-seq reference atlas demonstrates biologically meaningful and reproducible patterns of spatial localization across samples. B) Spatial mapping of TAM subpopulations using the same atlas reveals distinct and replicable spatial organization within the tumor microenvironment. C) Average spatial neighborhood analysis, integrating the spatial localization of each annotated cell type, identifies distinct and reproducible cellular neighborhoods within the lung tumor microenvironment, further validating the reliability and biological accuracy of our scRNA-seq atlas and its annotation.

**Figure 6.**
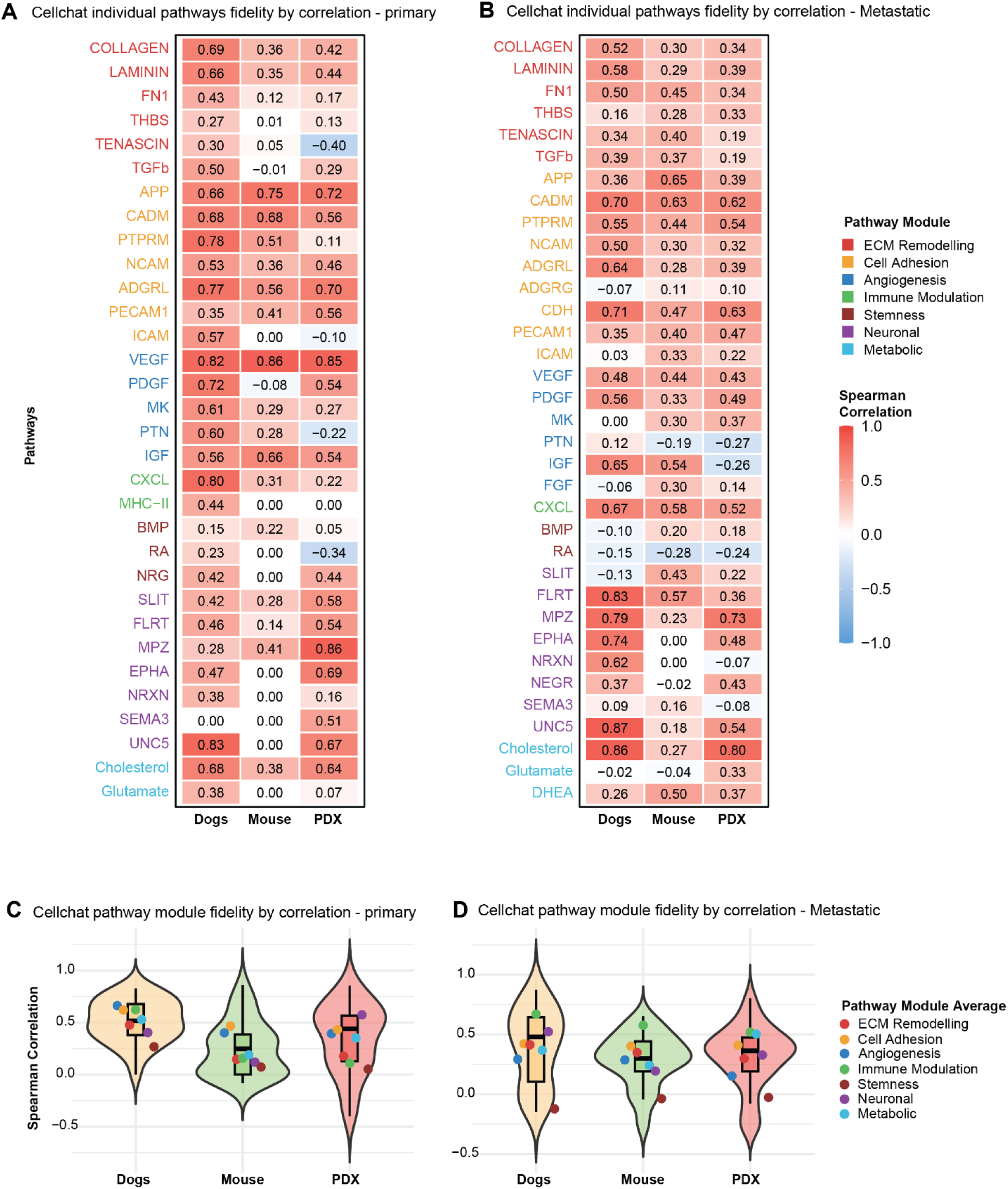
Cell–cell communication analysis reveals pathway-specific interspecies fidelity. CellChat-based cell–cell communication correlation analyses compare signaling pathway activity between dogs, mice, and PDX models relative to the human patient dataset at both primary and metastatic sites. Correlation analyses at the A) primary and B) metastatic sites show that the dog dataset most closely recapitulates human signaling interactions, whereas the mouse and PDX datasets exhibit pathway-specific similarities and divergences. Pathways are grouped and color-coded by functional modules, highlighting both conserved and model-specific communication programs across species. Violin plots showing the average activity of signaling pathway modules across species at the C) primary and D) metastatic sites. Each violin represents the distribution of module-level communication strengths aggregated across pathways within each functional category.

We found that the tumor subpopulations map onto three ordered regions. First, a leading-edge pattern emerged in which Fibrogenic and Interactive programs form a protective and expanding rim around the tumor core (Fig. 5A and Supplementary Fig. 12A). This region contains ECM– and cytokine-rich programs that may contribute to the reprogramming of host cells and the restructuring of the underlying matrix to create a more tumor-permissive environment as the lesion expands. Second, Proliferative, MP progenitor, and Basal progenitor programs were concentrated within the tumor core, indicating that maintenance and growth states are spatially protected in central regions rather than uniformly distributed (Fig. 5A and Supplementary Fig. 12A). Third, we observed a matrix-rich compartment marked by chondroid–osteoid deposition, stress, and hypoxia pathways (Fig. 5A and Supplementary Fig. 12A). The COMA subpopulation localized almost exclusively to this region, suggesting that the phenotype of these cells is defined by their confinement within a dense extracellular matrix.

The stromal and immune compartments consistently displayed a similar level of spatial organization. Fibrogenic TAMs and SCAR-associated TAMs were enriched at the tumor–stroma boundary, often co-localizing with fibrogenic and interactive tumor cells (Fig. 5B and Supplementary Fig. 12B), suggesting a coordinated role in sustaining the fibrotic and immunosuppressive niche. In contrast, osteoclast-like TAMs were found in discrete intra-tumoral pockets (Fig. 5B and Supplementary Fig. 12B), consistent with their function in osteoid degradation and matrix remodeling.

To add quantitative rigor to this visual assessment, we performed correlation analysis to identify distinct and recurrent spatial associations. This approach confirmed neighborhood structures that were highly consistent across all samples, indicating that the observed patterns reflect genuine biological organization rather than random spatial co-occurrence. We defined four distinct cellular neighborhoods-Tumor, Normal Alveoli, Tumor Periphery, and Immune compartments (Fig. 5C and Supplementary Fig. 13). Together, these findings demonstrate that osteosarcoma lung metastases harbor stable, functionally distinct spatial niches shaped by specific malignant, stromal, and immune programs, motivating further investigation into cell–cell crosstalk between malignant and stromal components across models and disease contexts.

### Cell-cell communication analysis reveals biologically relevant inter-species pathway-specific fidelity

We used the CellChat (46) algorithm to reconstruct potential tumor-host signaling networks based on ligand–receptor interactions across major cell types within each model and disease site. Using the CellChat-derived signaling matrices, we next performed a fidelity assessment to quantitatively evaluate how well PDX and mouse laboratory models or canine osteosarcoma recapitulate the network signaling dynamics of the human disease. We computed correlation using ligand-receptor signaling pathway composition, cell types receiving the signals, the number of interactions, and their strength for each dataset, compared to the human standard (Supplementary Fig. 14A). Pathways were divided into functional modules.

Correlation analysis of primary osteosarcoma suggests that, at a high level, canine tumors most closely recapitulated the cell-cell communication present within human tumors, especially for pathways involved in angiogenesis, cell adhesion, ECM remodeling, and immune modulation (Figs. 6A, 6C and Supplementary Fig. 14B). Murine models best preserved networks associated with angiogenesis and cell adhesion, but poorly reflected stemness pathways and several ECM ligand-receptor networks (Fig. 6A, 6C). PDXs largely retained angiogenesis, cell adhesion, neuronal, and metabolic modules, but showed the lowest fidelity for immune modulation and select ECM/angiogenic pathways, consistent with their immunodeficient nature (Figs. 6A, 6C).

In the metastatic lesions, canine tumors maintained the highest fidelity to human cell-cell communication in neuronal, metabolic, immune, angiogenesis, and cell adhesion modules (Fig. 6B, 6D, and Supplementary Fig. 14C), with stemness and some ECM components exhibiting lower fidelity (Figs. 6B, 6D). Murine metastases best preserved immune modulation, cell adhesion, and select ECM and angiogenic pathways, but showed lower fidelity for stemness and several ECM and neuronal components (Figs. 6B, 6D). PDXs largely retained metabolic, cell adhesion, and neuronal programs, with lower fidelity for stemness and select angiogenic and neuronal modules (Figs. 6B, 6D). Overall, dogs most comprehensively model human signaling, murine models show intermediate fidelity with select pathway preservation, and PDXs reflect their immunodeficient and partial ECM/angiogenic representation.

### Tumor–host communication through FN1 remodels the lung epithelium into a permissive metastatic niche

Building on this ligand–receptor–based communication framework, we systematically examined whether specific malignant cell states engage lung cell types with reprogramming signals that remodel the lung microenvironment. To quantify signaling differences, we computed average fold-change enrichment across pre-selected CellChat pathways, comparing human lung metastases to primary tumors. Surprisingly, the lung metastases displayed marked enrichment of extracellular matrix ECM–related signaling when compared to primary bone lesions with fibronectin (FN1) emerging as the ligand with the highest metastasis-specific normalized probability (Fig. 7A). This result was intriguing, given our prior report implicating FN1-dominant fibrogenesis as a process essential for metastatic colonization (78).

**Figure 7.**
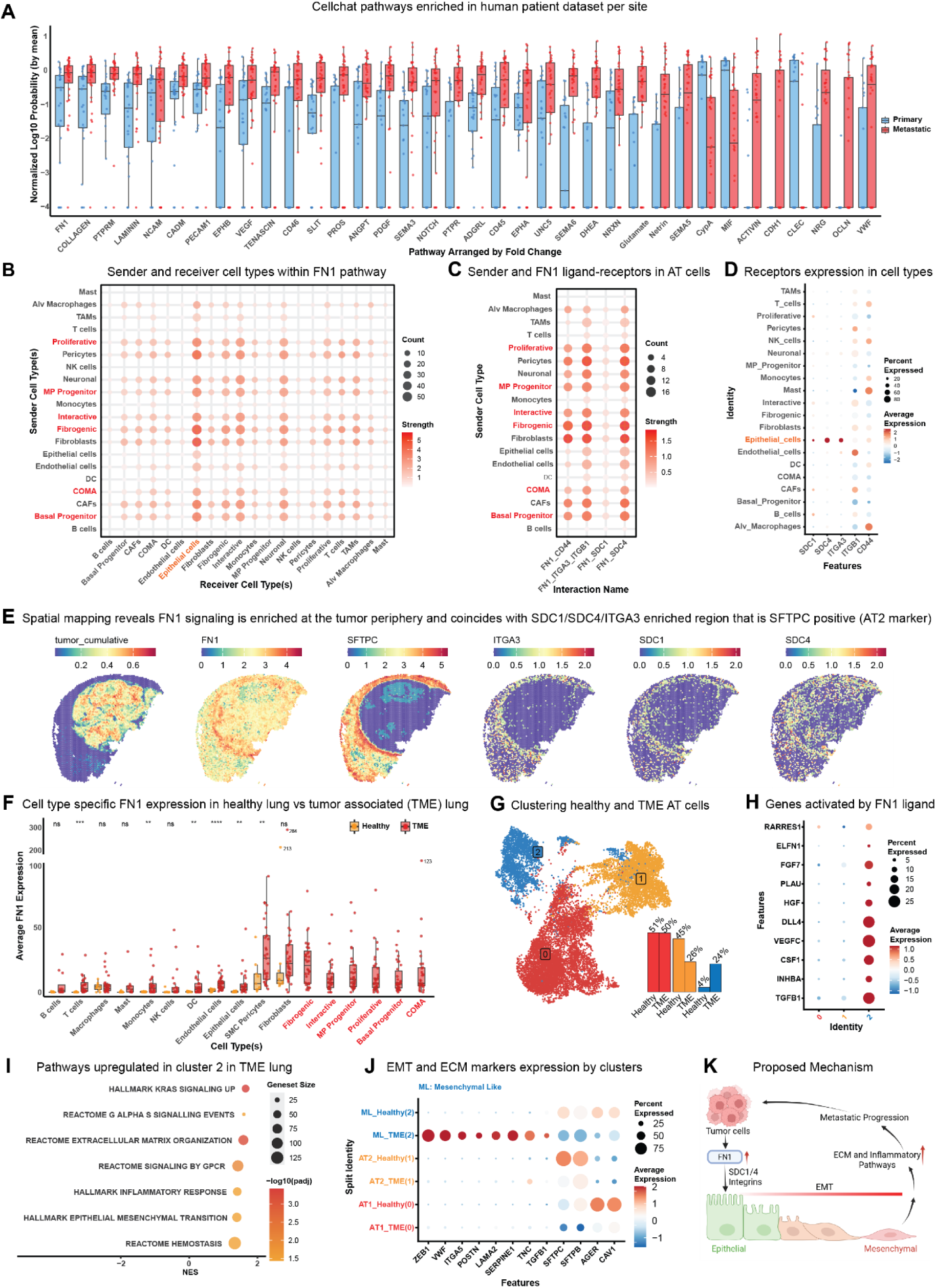
Tumor–host communication through FN1 remodels the lung epithelium into a permissive metastatic niche. A) Boxplot of cell–cell communication pathways significantly enriched in the human patient dataset at primary and metastatic sites. Pathways are normalized to the site with the highest average, with each dot representing an individual sample; FN1 ranks highest by normalized fold change. B) Dot plot showing the strength and frequency of FN1 signaling between sender and receiver cell types in the metastatic lung niche, revealing tumor and mesenchymal cells as primary senders and epithelial cells as top receiver. C) Dot plot highlighting FN1 ligand sender cell types and their corresponding receptor pairs in epithelial cells, revealing SDC1, SDC4, ITGA3, ITGA1, and CD44 as receptors. D) Dot plot of receptor expression for FN1 across cell types in the metastatic niche, showing receptor expression for SDC1, SDC4, and ITGA3 is largely restricted to epithelial cells. E) Spatial visualization of cellchat-inferred fn1 signaling in the visium dataset shows that fn1 ligand activity is concentrated along the outer tumor margin. the same regions are enriched for sftpc positive alveolar AT2 cells co-expressing the fn1 receptors sdc1, sdc4, and itga3, indicating localized tumor–epithelial communication at the invasive front. F) Cell-type–specific comparison of FN1 expression in healthy versus tumor-associated lung tissue, revealing elevated FN1 levels in the tumor microenvironment, primarily contributed by tumor and mesenchymal cells. G) Clustering of healthy and tumor-associated epithelial cell subsets, with an inset bar plot showing the proportion of cells from each dataset in each cluster; cluster 2 is enriched for tumor-associated epithelial cells. H) Dot plot showing expression of FN1-activated genes informed from the Nichenet ligand-target dataset, mapping exclusively to cluster 2 in the tumor-associated dataset. I) GSEA analysis with epithelial cells from the tumor-associated dataset, revealing that cluster 2 exhibits upregulated EMT, inflammatory, and ECM-related programs relative to other clusters. J) Dot plot split by cluster or cell type, and dataset demonstrating that cluster 2 from the tumor tumor-associated dataset shows upregulated EMT and ECM markers with concomitant downregulation of AT1 and AT2 markers. K) Proposed model of FN1-driven epithelial reprogramming in the metastatic lung niche.

To elucidate FN1-derived signaling interactions that might facilitate malignant progression, we examined the cell-specific components driving the identification of this ligand-receptor signature. In this analysis, epithelial cells were the strongest responders to the FN1-mediated signal, while tumor and tumor-associated mesenchymal cells were identified as primary sources of the signal (Fig. 7B). Closer inspection revealed that multiple epithelial cell-specific interactions could be initiated by FN1 signaling, with potential activation of receptors such as SDC1, SDC4, CD44 and key integrins (Fig. 7C). Notably, SDC1, SDC4, and ITGA3 were almost exclusively expressed within epithelial cells, supporting the idea that these may be key receptor components of this axis (Fig. 7D). Spatial mapping of cell-type markers and ligand–receptor pairs revealed a consistent pattern in which cells at the tumor-lung interface show elevated FN1 expression, while adjacent epithelial cells exhibit upregulation of the corresponding receptors (Fig. 7E). At the level of whole samples, FN1 expression was markedly upregulated in tumor subpopulations and several tumor-associated stromal cell types, when compared to their counterparts growing within healthy lung (Fig. 7F).

To determine whether these FN1-driven interactions were associated with a distinct epithelial response in the metastatic niche, we isolated alveolar epithelial cells from both healthy and tumor-associated lung samples (Fig. 7G). Following reprocessing and Harmony integration, three distinct epithelial clusters were identified. Among them, cluster 2 showed marked enrichment within the tumor-associated lung datasets, suggesting the emergence of a tumor-programmed state (Fig. 7G). To determine whether this reprogrammed state represented a response to FN1 signaling, we leveraged the NicheNet (45) ligand–target database to examine the expression of downstream FN1-responsive genes, including TGFB1, INHBA, CSF1, VEGFC, DLL4, HGF, PLAU, FGF7, ELFN1, and RARRES1 (Fig. 7H). Cluster 2 showed high expression of these genes downstream of FN1, supporting the hypothesis that these cells do indeed exhibit a functional response to FN1-dependent signals (Fig. 7H).

Finally, we performed a GSEA comparing cluster 2 with the remaining epithelial clusters. This analysis revealed significant upregulation of KRAS signaling, ECM organization, inflammatory response, and epithelial-to-mesenchymal transition (EMT) pathways in cluster 2 (Fig. 7I). KRAS upregulation has been reported to be connected to the exit from the epithelial state and the acquisition of a mesenchymal phenotype (79–81). Consistent with this, ECM– and EMT-associated gene signatures, such as ZEB1 (80,82), VWF (80), ITGA5 (80), POSTN (83), LAMA2 (80),

SERPINE1 (84), and TNC (85), were exclusively elevated in cluster 2, whereas cluster 0 expressed canonical AT1 markers and cluster 1 expressed AT2 markers (Fig. 7J). Interestingly, cluster 2 co-expressed both AT1 and AT2 alveolar epithelial markers at reduced levels compared to the other clusters, indicating a transitional epithelial state undergoing partial EMT (Fig. 7J). Together, these data indicate that fibrogenic and interactive tumor cells at the tumor–host interface produce FN1, which acts on epithelial cells to induce an FN1-dependent epithelial-to-mesenchymal–like transition, a phenotype strongly associated with pathogenic fibrosis in other diseases (11–13) (Fig. 7K).

## Discussion

Osteosarcoma is a highly heterogeneous malignancy, exhibiting both extensive intratumoral diversity and substantial inter-patient variation. Despite well-documented molecular complexity, it remains unclear how these tumors self-organize and adapt to metastatic sites, limiting our understanding of disease progression and therapeutic response (86,87). By integrating, aligning, and consistently annotating a comprehensive scRNA-seq dataset comprising over 1 million cells/nuclei across human, canine, xenograft, and murine models, our study defines the conserved transcriptional and cellular architecture underlying the intratumoral complexity of osteosarcoma within primary bone and lung metastatic microenvironments. The resulting datasets provide a molecular reference that resolves both tumor– and stroma-intrinsic variation-and that benchmarks pathway-specific research model fidelity-with unprecedented resolution.

A key innovation that enhances the rigor of this work is the cross-species truth set pipeline, which leverages properties of the multi-species dataset to overcome the objective limitations of CNV-based tools for tumor identification. Conventional algorithms such as SCEVAN (21), scATOMIC (22), and CopyKAT (23) produced highly inconsistent results in this genomically complex tumor type, with interrater agreement rates only slightly better than chance. These methods were particularly error-prone in samples from metastatic sites, likely due to the diverse cellular milieu within the lung. In contrast, our truth set–anchored classifier, validated against both synthetic mixtures and orthogonal SNV-based approaches (ScanBit), achieved over 95% accuracy and a very high degree of interrater consistency. This rigorous approach to the particularly challenging problem of reliably identifying tumor cells could be broadly applicable to other heterogeneous malignancies, helping to address a key challenge in annotation.

Within the tumor compartment, we identified six conserved subpopulations spanning proliferative, matrix-producing, progenitor-like, and inflammatory phenotypes. These states not only recur across species but also coexist within individual tumors, underscoring the profound intratumoral heterogeneity that characterizes osteosarcoma. RNA velocity and regulon activity analyses revealed a putative lineage hierarchy in which progenitor and proliferative states give rise to fibrogenic, COMA, and inflammatory phenotypes. The regulon landscape further supports these findings, with progenitor populations enriched for regulators such as E2F1, NFIA/NFIC, and PBX1, whereas ZEB1, PRDM1, and JUN dominate differentiated states.

It is interesting to consider how our results extend and complement the recent reports of Lopez-Fuentes et al (88). While our single-cell analyses were designed to resolve intra-tumoral heterogeneity, their bulk epigenetic profiling identified inter-patient variability and defined two major osteosarcoma subtypes—EOD and LOD—driven by RUNX2 or FOS/JUN activity, respectively, with distinct phenotypic and clinical features. Although most tumors in their study were classified as predominantly EOD or LOD, a subset exhibited mixed epigenetic states. In contrast, our analysis consistently identifies multiple tumor cell subpopulations coexisting within individual tumors, each associated with distinct but conserved regulatory programs. Notably, these subpopulation-specific regulons align with the same core transcriptional axes described in their work, suggesting that the bulk-defined subtypes may reflect shifts in the relative abundance of underlying cellular states rather than entirely distinct tumor classes. Extending this framework, our comparison of primary and metastatic samples further reveals a systematic shift in regulatory programs, with primary tumors enriched for stress-response and AP1–associated activity, consistent with an LOD-like phenotype, whereas metastatic lesions show attenuation of these programs alongside increased activation of developmental and chromatin-associated regulators, consistent with a more EOD-like state. Despite potential underestimation of enhancer-driven transcription factors by SCENIC, these observations support a model in which osteosarcoma tumors exist along a continuum of regulatory states, with metastatic progression favoring stabilization of EOD-like programs and a coordinated shift from stress– and inflammation-associated transcriptional programs toward more stable developmental and chromatin-associated regulatory states.

The interactive tumor subpopulation provides a potential mechanistic link between transcriptional plasticity and microenvironmental adaptation. Despite expression of immune-associated genes, these cells were confirmed to be *bona fide* tumor cells in PDX datasets, ruling out stromal contamination or doublet artifacts. Their enrichment in inflammatory and NF-κB regulons suggests an autocrine–paracrine axis enabling immune evasion and niche remodeling, reminiscent of the inflammation-coupled invasive states described in epithelial cancers (89–91). This dual phenotype could underlie the metastatic competence and therapeutic resistance observed in advanced disease.

In parallel, our cross-species stromal analysis revealed a conserved yet site-specifically remodeled immune microenvironment. TAM subsets, including inflammatory, IFN-responsive, fibrogenic, osteoclast-like, and scar-associated macrophages, were recurrent across models, with differentiation trajectories tracing from progenitor-like macrophages toward specialized states. Metastatic lung niches were enriched for scar-associated and immunosuppressive TAMs, implicating these subsets in metastatic colonization. Lymphoid, endothelial, and fibroblast populations were also conserved in canine and syngeneic mouse models, underscoring the complementary strengths of these systems for modeling tumor–immune dynamics.

The presence of osteoclast-like TAMs in the lung is interesting, as they are typically restricted to bone-resorptive niches rather than pulmonary tissue. Their emergence in metastatic lesions suggests that tumor–myeloid interactions may drive macrophages to adopt osteoclast-like programs outside the skeletal environment, highlighting an unexpected degree of microenvironmental plasticity and raising new questions about how osteosarcoma remodels distant metastatic sites.

Spatial deconvolution confirmed that transcriptional states map consistently into tissue-level organizational patterns, forming distinct neighborhoods. For example, fibrogenic and interactive tumor clusters co-localize with scar-associated TAMs and vascular stroma, forming micro-niches around the periphery of lesions that likely support metastatic outgrowth via inflammation and ECM production. This localization pattern aligns with the known biology of osteosarcoma, wherein interactive tumor cells actively engage with non-malignant tissues to modulate the surrounding microenvironment. Notably, this finding is highly consistent with previous work describing osteosarcoma as a fibrotic and wound-healing-like malignancy, whose progression depends on fibrosis-associated signaling and niche remodeling (78,92,93). Our integrative framework links cellular identity, spatial organization, and evolutionary trajectory, providing a systems-level view of osteosarcoma architecture.

We were surprised to find that intra-tumoral heterogeneity changed little from primary to metastatic lesions, a pattern observed across all species and models. However, while subsets of tumor cells within lesions showed minimal change, networks of cell-cell interactions differed significantly between primary and metastatic sites. Perhaps more surprisingly, lung lesions were characterized by cell-cell interactions that were dominated by extracellular matrix ligands, including collagen and fibronectin (FN1). As a functional consequence of this signaling, our findings suggest that tumor-derived FN1 engages syndecans (SDC1/SDC4) and integrins (e.g., ITGA3–ITGB1) on alveolar epithelial cells, with transcriptional patterns in the epithelial cells suggesting activation of downstream signaling. This signaling reprograms those epithelial cells into a hybrid inflammatory and EMT/profibrotic state that aids in building a fibrotic metastatic niche permissive for tumor lung colonization and metastatic progression. This is consistent with previous findings from our lab, where osteosarcoma cells upregulated FN1 expression when co-cultured with the epithelial cells, driving a wound-healing phenotype in epithelial cells (*in vitro* and *in vivo*) (78). Our findings are also consistent with others where syndecans and integrin receptors have been associated with a poor prognosis in diseases such as pulmonary fibrosis, lung adenocarcinoma, and osteosarcoma (11,94–96).

These findings, grounded in a cross-species dataset, reveal that osteosarcoma is shaped by conserved lineage hierarchies and adaptive stromal co-evolution rather than random cellular diversity. What appears chaotic at the end is the result of ordered and hierarchical tissue organization that follows constrained developmental pathways and anatomy. Tumor cells retain transcriptional heterogeneity across species and sites and are modulated by microenvironmental cues, adopting fibrogenic, interactive, and progenitor phenotypes. This framework identifies models best suited to study specific tumor-stroma interaction pathways. Our comparative analysis supports the idea that model fidelity is context dependent, with canine tumors showing strong concordance across anatomical sites while mouse and PDX models retaining clear value in many experimental settings. It further highlights FN1-mediated signaling as a key axis remodeling the lung epithelium into a pro-fibrotic, immune-permissive niche. By linking lineage programs, regulatory hierarchies, and spatial interactions, our study provides a unified model of osteosarcoma progression as a conserved failure of tissue repair and stromal adaptation, guiding mechanistic investigations and model selection.

## Acknowledgements

Funding for the generation of the single-cell and spatial datasets is provided in the original study publications. This reanalysis of those datasets was supported by an Alex’s Lemonade Stand Foundation Crazy 8 Award (RDR, RD, BEG), NIH R01CA260178 (RDR), NIH K08CA286845 (AGP), NIH K01OD028268 (HLG), NIH R37CA303990 (DPR),

Additional support was provided by a CancerFree KIDS Foundation Idea Award (RDR), the Boettcher Foundation Webb–Waring Biomedical Research Award (DPR), MIB Agents (RDR and DPR), and the American Kennel Club Canine Health Foundation #03015 (JFM). AGP is a Damon Runyon–Sohn Pediatric Cancer Fellow supported by the Damon Runyon Cancer Research Foundation and The Sohn Conference Foundation (DRSG-33P-20) and is also supported by the American Lebanese Syrian Associated Charities. JFM is supported by the Alvin and June Perlman Chair in Animal Oncology.

This research was supported, in part, by the Intramural Research Program of the National Institutes of Health (ZIABC012056 to TAM) and the NCI Childhood Cancer Data Initiative (3P30CA008748-54S3). The contributions of the NIH author(s) were made as part of their official duties as NIH federal employees, are in compliance with agency policy requirements, and are considered Works of the United States Government. However, the findings and conclusions presented in this paper are those of the author(s) and do not necessarily reflect the views of the NIH or the U.S. Department of Health and Human Services.

We would like to thank Mitzi Lewellen for providing support in maintaining records and databases, patient metadata, and sample follow-up at the University of Minnesota.

## References

1. Rajan S, Franz EM, McAloney CA, Vetter TA, Cam M, Gross AC, et al. Osteosarcoma tumors maintain intra-tumoral transcriptional heterogeneity during bone and lung colonization. BMC Biol 2023;21:98

2. Thomas DD, Lacinski RA, Lindsey BA. Single-cell RNA-seq reveals intratumoral heterogeneity in osteosarcoma patients: A review. J Bone Oncol 2023;39:100475

3. Wang D, Niu X, Wang Z, Song CL, Huang Z, Chen KN, et al. Multiregion Sequencing Reveals the Genetic Heterogeneity and Evolutionary History of Osteosarcoma and Matched Pulmonary Metastases. Cancer Res 2019;79:7–20

4. Rajan S, Zaccaria S, Cannon MV, Cam M, Gross AC, Raphael BJ, et al. Structurally Complex Osteosarcoma Genomes Exhibit Limited Heterogeneity within Individual Tumors and across Evolutionary Time. Cancer Res Commun 2023;3:564–75

5. Church AJ, Corson LB, Kao PC, Imamovic-Tuco A, Reidy D, Doan D, et al. Molecular profiling identifies targeted therapy opportunities in pediatric solid cancer. Nat Med 2022;28:1581–9

6. Kinnaman MD, Zaccaria S, Makohon-Moore A, Arnold B, Levine MF, Gundem G, et al. Subclonal Somatic Copy-Number Alterations Emerge and Dominate in Recurrent Osteosarcoma. Cancer Res 2023;83:3796–812

7. Zhou Y, Yang D, Yang Q, Lv X, Huang W, Zhou Z, et al. Single-cell RNA landscape of intratumoral heterogeneity and immunosuppressive microenvironment in advanced osteosarcoma. Nat Commun 2020;11:6322

8. Zheng X, Liu X, Zhang X, Zhao Z, Wu W, Yu S. A single-cell and spatially resolved atlas of human osteosarcomas. J Hematol Oncol 2024;17:71

9. He D, Che X, Zhang H, Guo J, Cai L, Li J, et al. Integrated single-cell analysis reveals heterogeneity and therapeutic insights in osteosarcoma. Discov Oncol 2024;15:669

10. Zheng X, Yuan J, Jin P, Wang X, Xu R, Feng H, et al. Single-Cell and Spatial Profiling of Tumor Microenvironment Heterogeneity in Human Osteosarcomas. bioRxiv 2025

11. Parimon T, Yao C, Habiel DM, Ge L, Bora SA, Brauer R, et al. Syndecan-1 promotes lung fibrosis by regulating epithelial reprogramming through extracellular vesicles. JCI Insight 2019;5

12. Griggs LA, Hassan NT, Malik RS, Griffin BP, Martinez BA, Elmore LW, et al. Fibronectin fibrils regulate TGF-beta1-induced Epithelial-Mesenchymal Transition. Matrix Biol 2017;60-61:157–75

13. Guerrero-Barbera G, Burday N, Costell M. Shaping Oncogenic Microenvironments: Contribution of Fibronectin. Front Cell Dev Biol 2024;12:1363004

14. Ammons DT, Hopkins LS, Cronise KE, Kurihara J, Regan DP, Dow S. Single-cell RNA sequencing reveals the cellular and molecular heterogeneity of treatment-naive primary osteosarcoma in dogs. Commun Biol 2024;7:496

15. Eigenbrood J, Wong N, Mallory P, Pereira JS, Williams D, Morris-Ii DW, et al. Spatial Profiling Identifies Regionally Distinct Microenvironments and Targetable Immunosuppressive Mechanisms in Pediatric Osteosarcoma Pulmonary Metastases. Cancer Res 2025;85:2320–37

16. Husted C, Seidel K, Karagiannis TT, Peterson C, Megquier K, Genereux D, et al. Multi-omic Longitudinal Analysis of Canine Osteosarcoma Identifies Inter-Patient Heterogeneity and Immune Enrichment in Metastatic Lesions. bioRxiv 2026:2026.01.05.696411

17. Kinsella RJ, Kahari A, Haider S, Zamora J, Proctor G, Spudich G, et al. Ensembl BioMarts: a hub for data retrieval across taxonomic space. Database (Oxford) 2011;2011:bar030

18. Hao Y, Hao S, Andersen-Nissen E, Mauck WM, 3rd, Zheng S, Butler A, et al. Integrated analysis of multimodal single-cell data. Cell 2021;184:3573–87 e29

19. Zappia L, Oshlack A. Clustering trees: a visualization for evaluating clusterings at multiple resolutions. Gigascience 2018;7

20. Korsunsky I, Millard N, Fan J, Slowikowski K, Zhang F, Wei K, et al. Fast, sensitive and accurate integration of single-cell data with Harmony. Nat Methods 2019;16:1289–96

21. De Falco A, Caruso F, Su XD, Iavarone A, Ceccarelli M. A variational algorithm to detect the clonal copy number substructure of tumors from scRNA-seq data. Nat Commun 2023;14:1074

22. Nofech-Mozes I, Soave D, Awadalla P, Abelson S. Pan-cancer classification of single cells in the tumour microenvironment. Nature Communications 2023;14

23. Gao R, Bai S, Henderson YC, Lin Y, Schalck A, Yan Y, et al. Delineating copy number and clonal substructure in human tumors from single-cell transcriptomes. Nat Biotechnol 2021;39:599–608

24. Cannon M, Gust M, Gross A, Cam M, Reinecke J, Garcia LJ, et al. SCANBIT facilitates identification of tumor cell populations in scRNAseq data using pseudobulked SNV calls. bioRxiv 2026–01–28

25. Aran D, Looney AP, Liu L, Wu E, Fong V, Hsu A, et al. Reference-based analysis of lung single-cell sequencing reveals a transitional profibrotic macrophage. Nat Immunol 2019;20:163–72

26. Sikkema L, Ramirez-Suastegui C, Strobl DC, Gillett TE, Zappia L, Madissoon E, et al. An integrated cell atlas of the lung in health and disease. Nat Med 2023;29:1563–77

27. Aibar S, Gonzalez-Blas CB, Moerman T, Huynh-Thu VA, Imrichova H, Hulselmans G, et al. SCENIC: single-cell regulatory network inference and clustering. Nat Methods 2017;14:1083–6

28. Duval F, Lourenco J, Hicham M, Boivin G, Guichard A, Wyser-Rmili C, et al. Trajectories of macrophage ontogeny and reprogramming in cancer. iScience 2025;28:112498

29. Ma RY, Black A, Qian BZ. Macrophage diversity in cancer revisited in the era of single-cell omics. Trends Immunol 2022;43:546–63

30. Franzen O, Gan LM, Bjorkegren JLM. PanglaoDB: a web server for exploration of mouse and human single-cell RNA sequencing data. Database (Oxford) 2019;2019

31. Love MI, Huber W, Anders S. Moderated estimation of fold change and dispersion for RNA-seq data with DESeq2. Genome Biol 2014;15:550

32. Liberzon A, Subramanian A, Pinchback R, Thorvaldsdottir H, Tamayo P, Mesirov JP. Molecular signatures database (MSigDB) 3.0. Bioinformatics 2011;27:1739–40

33. Bergen V, Lange M, Peidli S, Wolf FA, Theis FJ. Generalizing RNA velocity to transient cell states through dynamical modeling. Nat Biotechnol 2020;38:1408–14

34. Hawkins AG, Shapiro JA, Spielman SJ, Mejia DS, Venkatesh Prasad D, Ichihara N, et al. The Single-cell Pediatric Cancer Atlas: Data portal and open-source tools for single-cell transcriptomics of pediatric tumors. bioRxiv 2025

35. La Manno G, Soldatov R, Zeisel A, Braun E, Hochgerner H, Petukhov V, et al. RNA velocity of single cells. Nature 2018;560:494–8

36. Wolf FA, Angerer P, Theis FJ. SCANPY: large-scale single-cell gene expression data analysis. Genome Biol 2018;19:15

37. Germain PL, Lun A, Garcia Meixide C, Macnair W, Robinson MD. Doublet identification in single-cell sequencing data using scDblFinder. F1000Res 2021;10:979

38. Lun ATL, Riesenfeld S, Andrews T, Dao TP, Gomes T, participants in the 1st Human Cell Atlas J, et al. EmptyDrops: distinguishing cells from empty droplets in droplet-based single-cell RNA sequencing data. Genome Biol 2019;20:63

39. Neavin D, Senabouth A, Arora H, Lee JTH, Ripoll-Cladellas A, sc-e QC, et al. Demuxafy: improvement in droplet assignment by integrating multiple single-cell demultiplexing and doublet detection methods. Genome Biol 2024;25:94

40. Gayoso A, Shor J. JonathanShor/DoubletDetection: doubletdetection v4.3.0.post1.

41. Germain P-L, Lun A, Garcia Meixide C, Macnair W, Robinson MD, Germain P-L, et al. Doublet identification in single-cell sequencing data using scDblFinder. 2022/05/16.

42. Bais AS, Kostka D. scds: computational annotation of doublets in single-cell RNA sequencing data. Bioinformatics 2020;36:1150–8

43. Cable DM, Murray E, Zou LS, Goeva A, Macosko EZ, Chen F, et al. Robust decomposition of cell type mixtures in spatial transcriptomics. Nat Biotechnol 2022;40:517–26

44. Lee S-I. Developing a bivariate spatial association measure: An integration of Pearson’s r and Moran’s I. Journal of Geographical Systems 2001;3:369–85

45. Browaeys R, Saelens W, Saeys Y. NicheNet: modeling intercellular communication by linking ligands to target genes. Nat Methods 2020;17:159–62

46. Jin S, Guerrero-Juarez CF, Zhang L, Chang I, Ramos R, Kuan CH, et al. Inference and analysis of cell-cell communication using CellChat. Nat Commun 2021;12:1088

47. Hao Y, Stuart T, Kowalski MH, Choudhary S, Hoffman P, Hartman A, et al. Dictionary learning for integrative, multimodal and scalable single-cell analysis. Nat Biotechnol 2024;42:293–304

48. Van de Sande B, Flerin C, Davie K, De Waegeneer M, Hulselmans G, Aibar S, et al. A scalable SCENIC workflow for single-cell gene regulatory network analysis. Nat Protoc 2020;15:2247–76

49. Avin KG, Dominguez JM, 2nd, Chen NX, Hato T, Myslinski JJ, Gao H, et al. Single-cell RNAseq provides insight into altered immune cell populations in human fracture nonunions. J Orthop Res 2023;41:1060–9

50. Angeles-Lopez QD, Rodriguez-Lopez J, Agudelo Garcia P, Calyeca J, Alvarez D, Bueno M, et al. Regulation of lung progenitor plasticity and repair by fatty acid oxidation. JCI Insight 2025;10

51. Korotkevich G, Sukhov V, Budin N, Shpak B, Artyomov MN, Sergushichev A. Fast gene set enrichment analysis. bioRxiv 2021

52. R E, EF W. AP-1: a double-edged sword in tumorigenesis – PubMed. Nature reviews Cancer 2003 Nov;3

53. Wortel IM, Meer LTvd, Kilberg MS, Leeuwen FNv. Surviving Stress: Modulation of ATF4-Mediated Stress Responses in Normal and Malignant Cells. Trends in endocrinology and metabolism: TEM 2017 Aug 7;28

54. JJ OS, R P. JAK and STAT signaling molecules in immunoregulation and immune-mediated disease – PubMed. Immunity 04/20/2012;36

55. Ren Q, Liu Z, Wu L, Yin G, Xie X, Kong W, et al. C/EBPβ: The structure, regulation, and its roles in inflammation-related diseases. Biomedicine & Pharmacotherapy 2023/12/31;169

56. Cai J, Chen J, Jiang Y, Liu B. Meis1 Negatively Regulates Epithelial-Mesenchymal Transition via Wnt/β-catenin Pathway in Oral Submucous Fibrosis. International Dental Journal 2026 Jan 23;76

57. Attwooll C, Denchi EL, Helin K, Attwooll C, Denchi EL, Helin K. The E2F family: specific functions and overlapping interests. The EMBO Journal 2004 23:24 2004–11–11;23

58. Ren B, Cam H, Takahashi Y, Volkert T, Terragni J, Young RA, et al. E2F integrates cell cycle progression with DNA repair, replication, and G2/M checkpoints. Genes & Development 2002–01–15;16

59. Li W-f, Herkilini A, Tang Y, Huang P, Song G-b, Miyagishi M, et al. The transcription factor PBX3 promotes tumor cell growth through transcriptional suppression of the tumor suppressor p53. Acta Pharmacologica Sinica 2021 Feb 1;42

60. Qu K, Mo S, Huang J, Liu S, Zhang S, Shen J, et al. SPI1-KLF1/LYL1 axis regulates lineage commitment during endothelial-to-hematopoietic transition from human pluripotent stem cells. iScience 2024 Jun 28;27

61. Garrett-Sinha LA. An update on the roles of transcription factor Ets1 in autoimmune diseases. WIREs mechanisms of disease 2023 Aug 10;15

62. Feng L, Zhang H, Liu T. Multifaceted roles of IKZF1 gene, perspectives from bench to bedside. Frontiers in Oncology 2024 Jun 24;14

63. Liu M, Xing Y, Tan J, Chen X, Xue Y, Qu L, et al. Frontiers | Comprehensive summary: the role of PBX1 in development and cancers. Frontiers in Cell and Developmental Biology 2024/07/26;12

64. Wang J, Zhang J, Xu L, Zheng Y, Ling D, Yang Z. Expression of HNF4G and its potential functions in lung cancer. Oncotarget 2017 Dec 4;9

65. Deen D, Butter F, Daniels DE, Ferrer-Vicens I, Ferguson DCJ, Holland ML, et al. Identification of the transcription factor MAZ as a regulator of erythropoiesis. Blood Advances 2021 Aug 5;5

66. He Y, Dupree J, Wang J, Sandoval J, Li J, Liu H, et al. The Transcription Factor Yin Yang 1 Is Essential for Oligodendrocyte Progenitor Differentiation. Neuron 2007/07/19;55

67. Liu Y, El-Naggar S, Darling DS, Higashi Y, Dean DC. ZEB1 Links Epithelial-Mesenchymal Transition and Cellular Senescence. Development (Cambridge, England) 2008 Feb;135

68. E S, M K. AP-1 as a regulator of cell life and death – PubMed. Nature cell biology 2002 May;4

69. Lau CM, Tiniakou I, Perez OA, Kirkling ME, Yap GS, Hock H, et al. Transcription factor Etv6 regulates functional differentiation of cross-presenting classical dendritic cells. The Journal of Experimental Medicine 2018 Sep 3;215

70. Nuclear Factor I A – an overview | ScienceDirect Topics. Journal of Molecular Biology 2024;436

71. HMGA2 – an overview | ScienceDirect Topics. Experimental Hematology 2007;35

72. Transcription Factor Ets 1 – an overview | ScienceDirect Topics. Free Radical Biology and Medicine 2004;37

73. Zhou Y, Huang T, Cheng ASL, Yu J, Kang W, To KF. The TEAD Family and Its Oncogenic Role in Promoting Tumorigenesis. International Journal of Molecular Sciences 2016 Jan 21;17

74. Di Pompo G, Mitsiadis TA, Pagella P, Pasquarelli A, Bettini G, Sabattini S, et al. Mesenchymal stroma drives axonogenesis and nerve-induced aggressiveness in osteosarcoma. J Exp Clin Cancer Res 2025;44:276

75. Liu B, Ma W, Jha RK, Gurung K. Cancer stem cells in osteosarcoma: recent progress and perspective. Acta Oncol 2011;50:1142–50

76. Jian C, Wang B, Mou H, Zhang Y, Yang C, Huang Q, et al. A GAD1 inhibitor suppresses osteosarcoma growth through the Wnt/beta-catenin signaling pathway. Heliyon 2024;10:e31444

77. Cervantes-Villagrana RD, Albores-Garcia D, Cervantes-Villagrana AR, Garcia-Acevez SJ. Tumor-induced neurogenesis and immune evasion as targets of innovative anti-cancer therapies. Signal Transduct Target Ther 2020;5:99

78. Reinecke JB, Jimenez Garcia L, Gross AC, Cam M, Cannon MV, Gust MJ, et al. Aberrant activation of wound healing programs within the metastatic niche facilitates lung colonization by osteosarcoma cells. Cold Spring Harbor Laboratory; 2024.

79. McFaline-Figueroa JL, Hill AJ, Qiu X, Jackson D, Shendure J, Trapnell C. A pooled single-cell genetic screen identifies regulatory checkpoints in the continuum of the epithelial-to-mesenchymal transition. Nat Genet 2019;51:1389–98

80. Malagoli Tagliazucchi G, Wiecek AJ, Withnell E, Secrier M. Genomic and microenvironmental heterogeneity shaping epithelial-to-mesenchymal trajectories in cancer. Nat Commun 2023;14:789

81. Rhim AD, Mirek ET, Aiello NM, Maitra A, Bailey JM, McAllister F, et al. EMT and dissemination precede pancreatic tumor formation. Cell 2012;148:349–61

82. Liu Y, El-Naggar S, Darling DS, Higashi Y, Dean DC. Zeb1 links epithelial-mesenchymal transition and cellular senescence. Development 2008;135:579–88

83. Shang Y, Liang Y, Zhang B, Wu W, Peng Y, Wang J, et al. Periostin-mediated activation of NF-kappaB signaling promotes tumor progression and chemoresistance in glioblastoma. Sci Rep 2025;15:13955

84. Ma Z, Li X, Yang S, Yang H, Zhang A, Li N, et al. Molecular Mechanisms of Neutrophil Extracellular Traps in Promoting Gastric Cancer Epithelial-Mesenchymal Transition Through SERPINE-1 Expression. J Biochem Mol Toxicol 2025;39:e70157

85. Kang X, Xu E, Wang X, Qian L, Yang Z, Yu H, et al. Tenascin-c knockdown suppresses vasculogenic mimicry of gastric cancer by inhibiting ERK-triggered EMT. Cell Death Dis 2021;12:890

86. Roberts RD, Lizardo MM, Reed DR, Hingorani P, Glover J, Allen-Rhoades W, et al. Provocative questions in osteosarcoma basic and translational biology: A report from the Children’s Oncology Group. Cancer 2019;125:3514–25

87. Odri GA, Tchicaya-Bouanga J, Yoon DJY, Modrowski D. Metastatic Progression of Osteosarcomas: A Review of Current Knowledge of Environmental versus Oncogenic Drivers. Cancers (Basel) 2022;14

88. Lopez-Fuentes E, Clugston AS, Lee AG, Sayles LC, Sorensen N, Pons Ventura MV, et al. Epigenetic and Transcriptional Programs Define Osteosarcoma Subtypes and Establish Targetable Vulnerabilities. Cancer Discov 2026;16:296–319

89. Huber MA, Azoitei N, Baumann B, Grunert S, Sommer A, Pehamberger H, et al. NF-kappaB is essential for epithelial-mesenchymal transition and metastasis in a model of breast cancer progression. J Clin Invest 2004;114:569–81

90. Oh A, Pardo M, Rodriguez A, Yu C, Nguyen L, Liang O, et al. NF-kappaB signaling in neoplastic transition from epithelial to mesenchymal phenotype. Cell Commun Signal 2023;21:291

91. Jin Y, Cai Q, Wang L, Ji J, Sun Y, Jiang J, et al. Paracrine activin B-NF-kappaB signaling shapes an inflammatory tumor microenvironment in gastric cancer via fibroblast reprogramming. J Exp Clin Cancer Res 2023;42:269

92. Reinecke JB, Jimenez Garcia L, Saraf AJ, Hinckley J, Gross AC, Le Pommellet H, et al. Metastasis-Initiating Osteosarcoma Subpopulations Establish Paracrine Interactions with Lung and Tumor Cells to Create a Metastatic Niche. Cancer Res 2025;85:4341–58

93. McAloney CA, Makkawi R, Budhathoki Y, Cannon MV, Franz EM, Gross AC, et al. Host-derived growth factors drive ERK phosphorylation and MCL1 expression to promote osteosarcoma cell survival during metastatic lung colonization. Cell Oncol (Dordr) 2024;47:259–82

94. Na KY, Bacchini P, Bertoni F, Kim YW, Park YK. Syndecan-4 and fibronectin in osteosarcoma. Pathology 2012;44:325–30

95. Toba-Ichihashi Y, Yamaoka T, Ohmori T, Ohba M. Up-regulation of Syndecan-4 contributes to TGF-beta1-induced epithelial to mesenchymal transition in lung adenocarcinoma A549 cells. Biochem Biophys Rep 2016;5:1–7

96. Zhu L, Xie L, Zhi Y, Huang Y, Chen H, Chen Z, et al. Fibrotic lung ECM upregulates SDC4/integrin-αvβ1 interaction and the interfering peptide SDC487-131 and its derivative peptides alleviate pulmonary fibrosis. Regenerative Biomaterials 2025

