## Supplementary material for "Integrative Single-cell and Spatial Transcriptomic Analysis of Osteosarcoma Reveals Conserved and Distinct Ecosystems Across Sites and Species": Supp Figures: CombinedSupplement.pdf

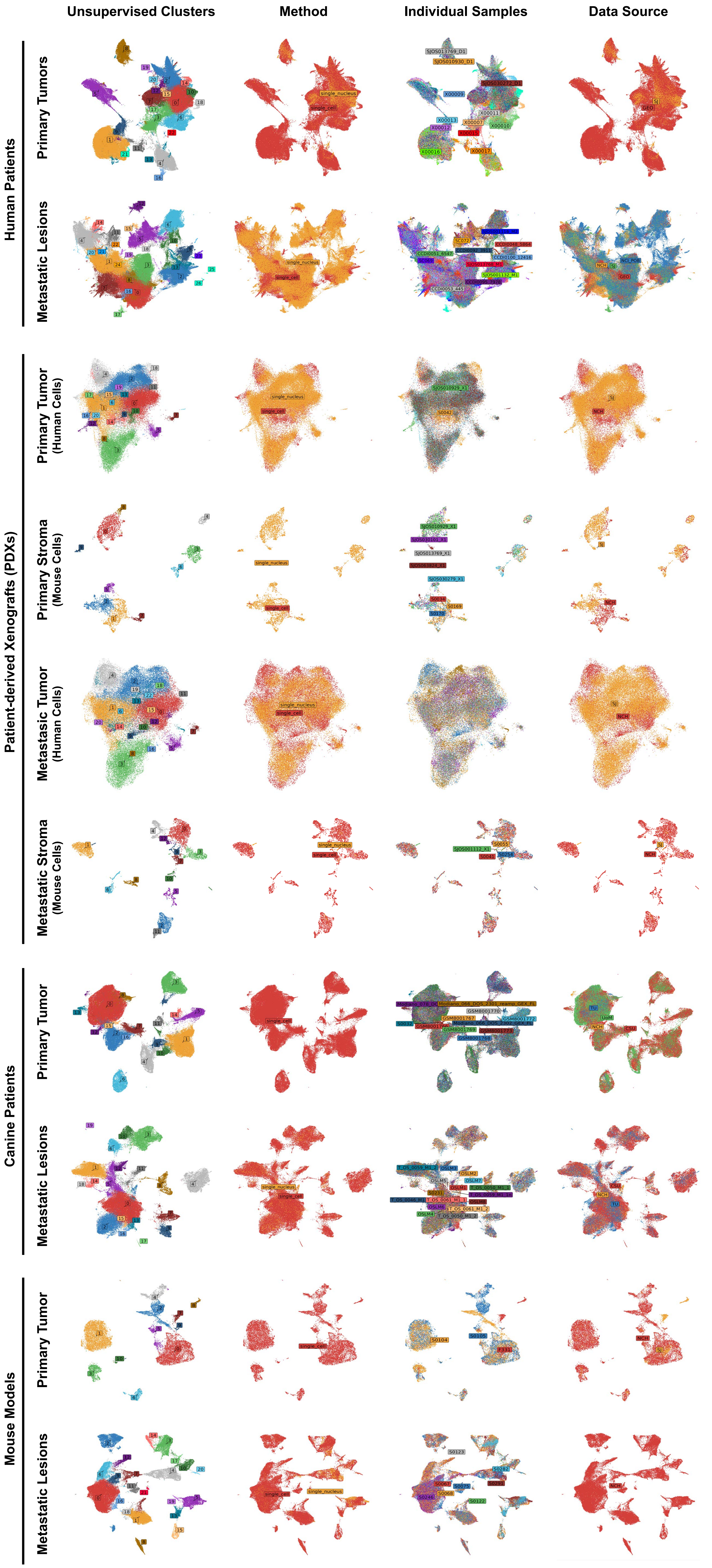


**Supplementary Figure 1 | Batch-associated variation is reduced after Harmony integration.** Dimensional reduction (DimPlot) visualizations across tumor and stromal compartments after Harmony integration are shown for (top to bottom) human patients, patient-derived xenografts, canine patients, and murine models of disease. Each is split by primary tumor (above) and metastatic lung lesions (below), reflecting integration like-by-like, given the distinct cell types present within bony and lung microenvironments. Each integrated dataset is colored by (left to right) unsupervised cluster assignment, capture method (single cell vs single nucleus), sample of origin, and site of origin. Following integration, cells show improved mixing across sequencing methods and data sources, suggesting effective correction of batch- and sample-associated systematic variation.

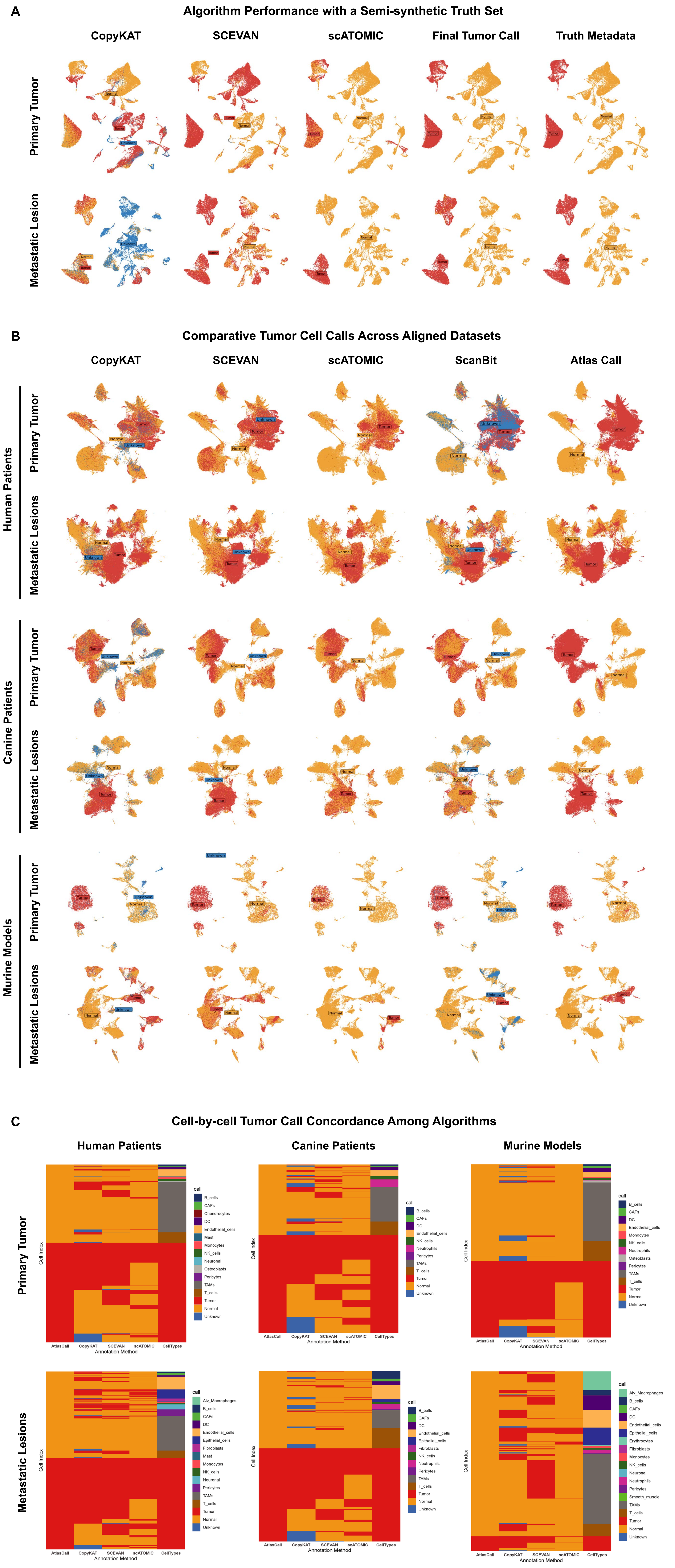


**Supplementary Figure 2 | Validation and performance of tumor cell identification algorithms.** A) Evaluation of algorithms using a semi-synthetic truth set, where tumor cell identity is known. Dimensional reduction (UMAP) plots compare CopyKAT, SCEVAN, scATOMIC, and our reference-based algorithm agains the known ground truth for a semi-synthetic primary tumor and metastatic lesion dataset. The reference-based algorithm demonstrates improved performance relative to established methods for tumor cell identification. ScanBit is not included in this analysis, as the nature of the semi-synthetic truth set would promote artifactually high accuracy. B) Comparison of tumor-normal classification algorithms in the harmonized real-world datasets. Dimensional reduction (UMAP) plots are again colored by the identity calls: tumor vs normal vs unknown. Of note, while data are shown, these conventional assessment methods may not apply to canine datasets, as conversion to human orthologues introduces artifacts in the determination of copy number variation (although gene A may be genomically close to gene B on an human chromosome, it is unlikely that this same proximity exists in the orthologous genes within the canine genome). Overall, ScanBit and the truth-set-based pipeline show strong concordance, whereas substantial discordance is noted in the results of the conventional algorithms. C) Cell-by-cell evaluation of tumor calls across methods. In these plots, each row is an individual cell and each column is a method for tumor cell identification. Stromal cell types are included to highlight cell populations frequently misclassified as tumor cells by established approaches. Tumor-associated macrophages (TAMs), endothelial, epithelial, and mesenchymal cells are commonly assigned as tumor cells by established methods.

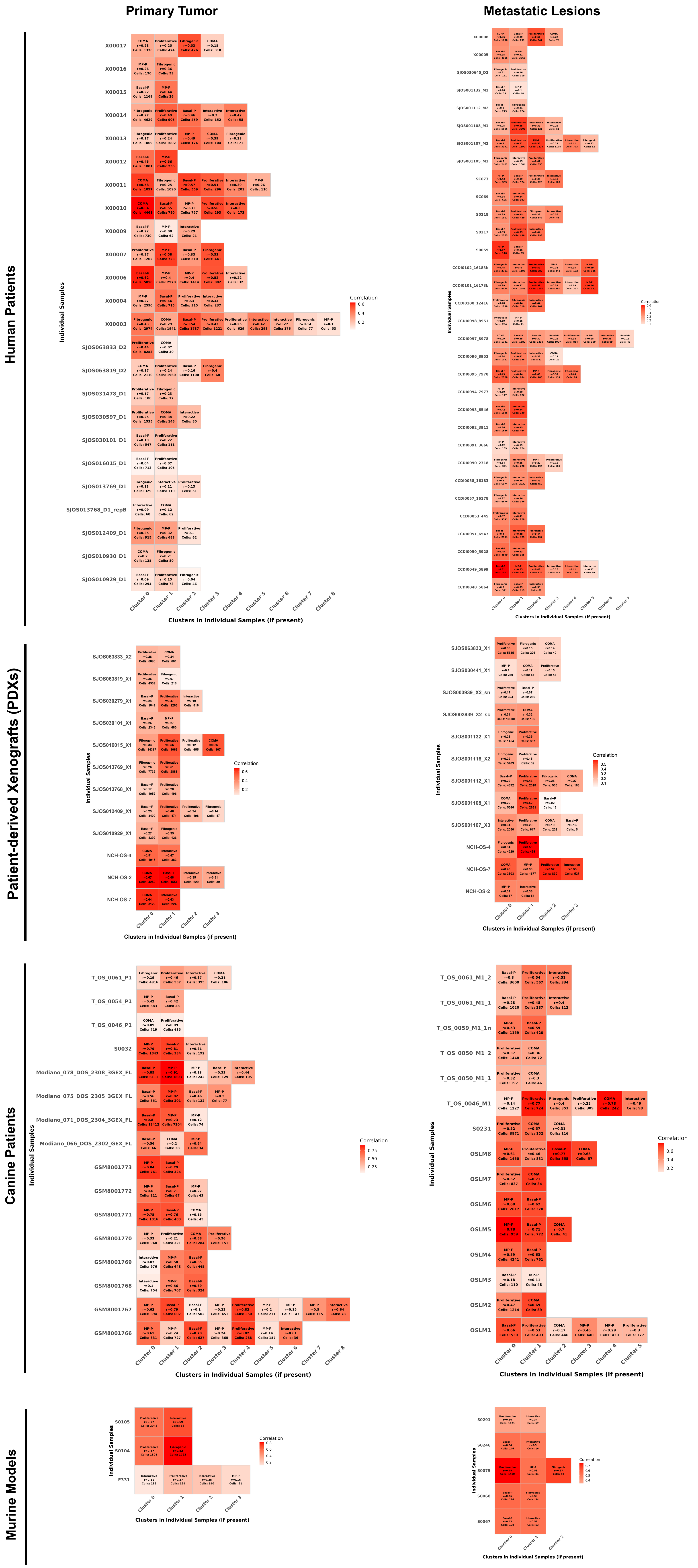


**Supplementary Figure 3 | Validation of Harmony integration across samples.** Tile heatmap showing validation of Harmony-based data integration. Tumor cells from each sample were reclustered independently, and correlation analysis was performed between clusters from individual samples and tumor subpopulations from the integrated dataset. The x-axis represents clusters from individual samples, and the y-axis represents individual samples. Heatmap values indicate correlations with clusters from the integrated Seurat object. High correlations between specific sample-level clusters and integrated clusters indicate that phenotypes identified in the integrated dataset are well represented within individual samples, supporting effective data integration.

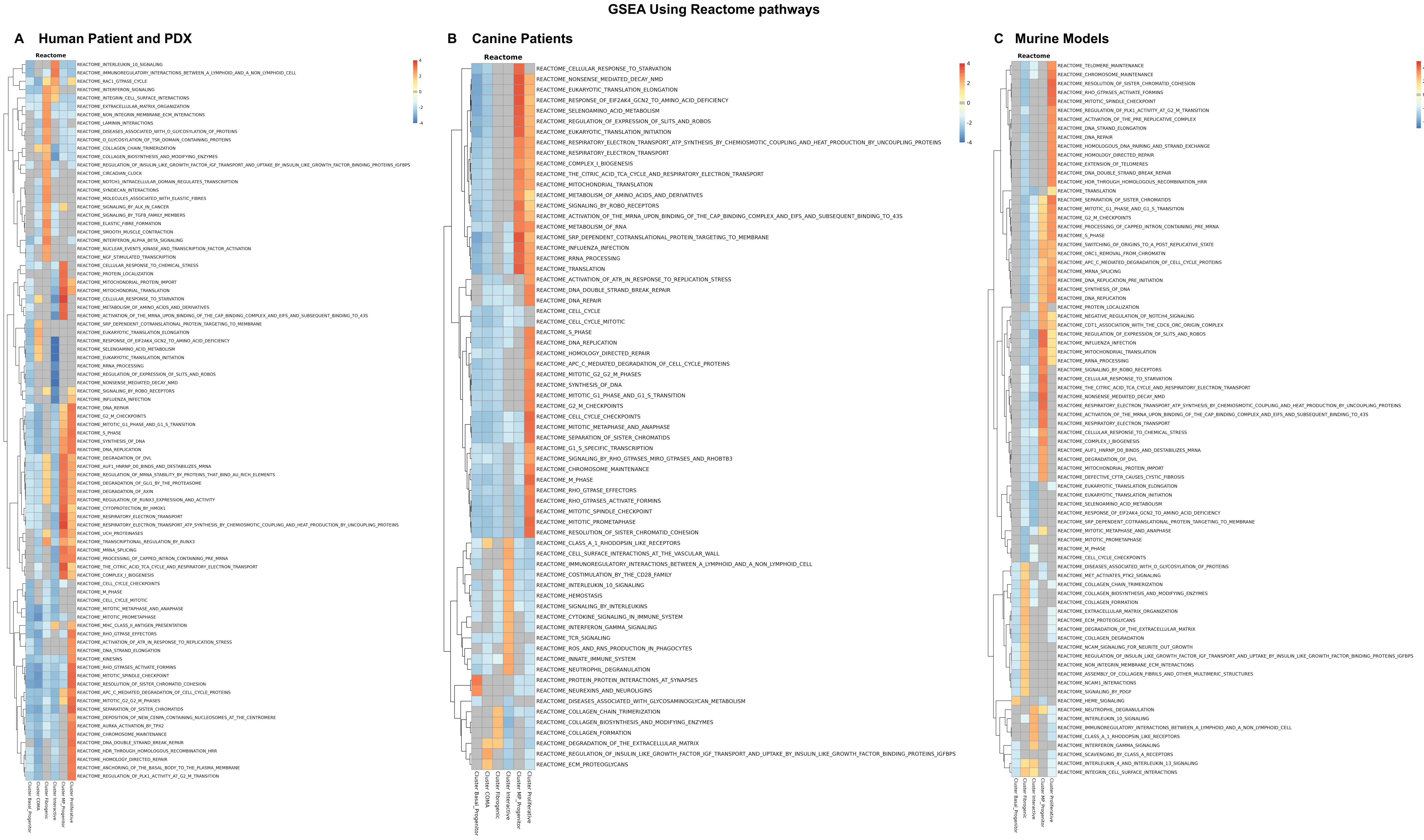


**Supplementary Figure 13 | Reactome pathway enrichment across tumor subpopulations in human, canine, and mouse tumors.** A) Reactome pathway gene set enrichment analysis (GSEA) performed across tumor subpopulations identified in human patient and patient-derived xenograft (PDX) tumor cells. Distinct transcriptional programs characterize each subpopulation: the basal progenitor subpopulation shows few or no significantly enriched pathways; the COMA subpopulation is enriched for stress- and starvation-associated pathways; the fibrogenic subpopulation shows enrichment of extracellular matrix (ECM)–related pathways; the interactive subpopulation is enriched for interleukin and inflammatory signaling pathways; the MP progenitor subpopulation shows enrichment of metabolic and oxidative phosphorylation pathways; and the proliferative subpopulation is enriched for cell-cycle–related pathways. B) Reactome pathway enrichment across canine tumor subpopulations following conversion of canine genes to human orthologs. Canine tumor subpopulations display pathway enrichment patterns that broadly mirror those observed in human tumors, supporting conservation of major transcriptional programs across species. C) Reactome pathway enrichment across mouse tumor subpopulations. While most mouse tumor subpopulations exhibit pathway enrichment patterns consistent with those observed in human and canine datasets, the COMA subpopulation detected in human and canine tumors is not identified in the mouse model, indicating partial but not complete cross-species conservation of tumor cell states.

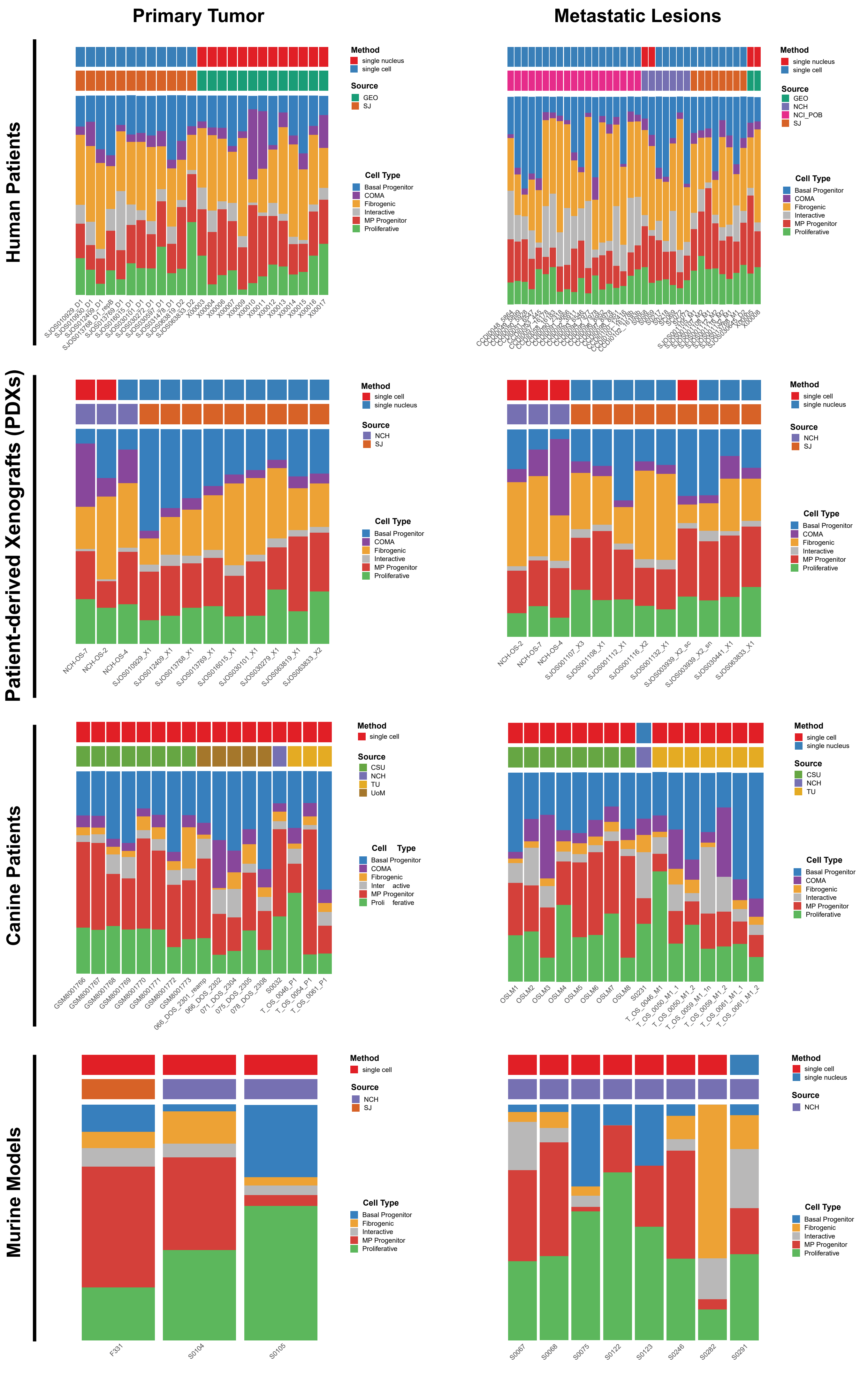


**Extended Data Figure 5 | Proportional composition of tumor subpopulations across samples and datasets.** Bar plots showing the proportional composition of tumor subpopulations across individual samples, sequencing modalities, and data sources for each species and tumor site.

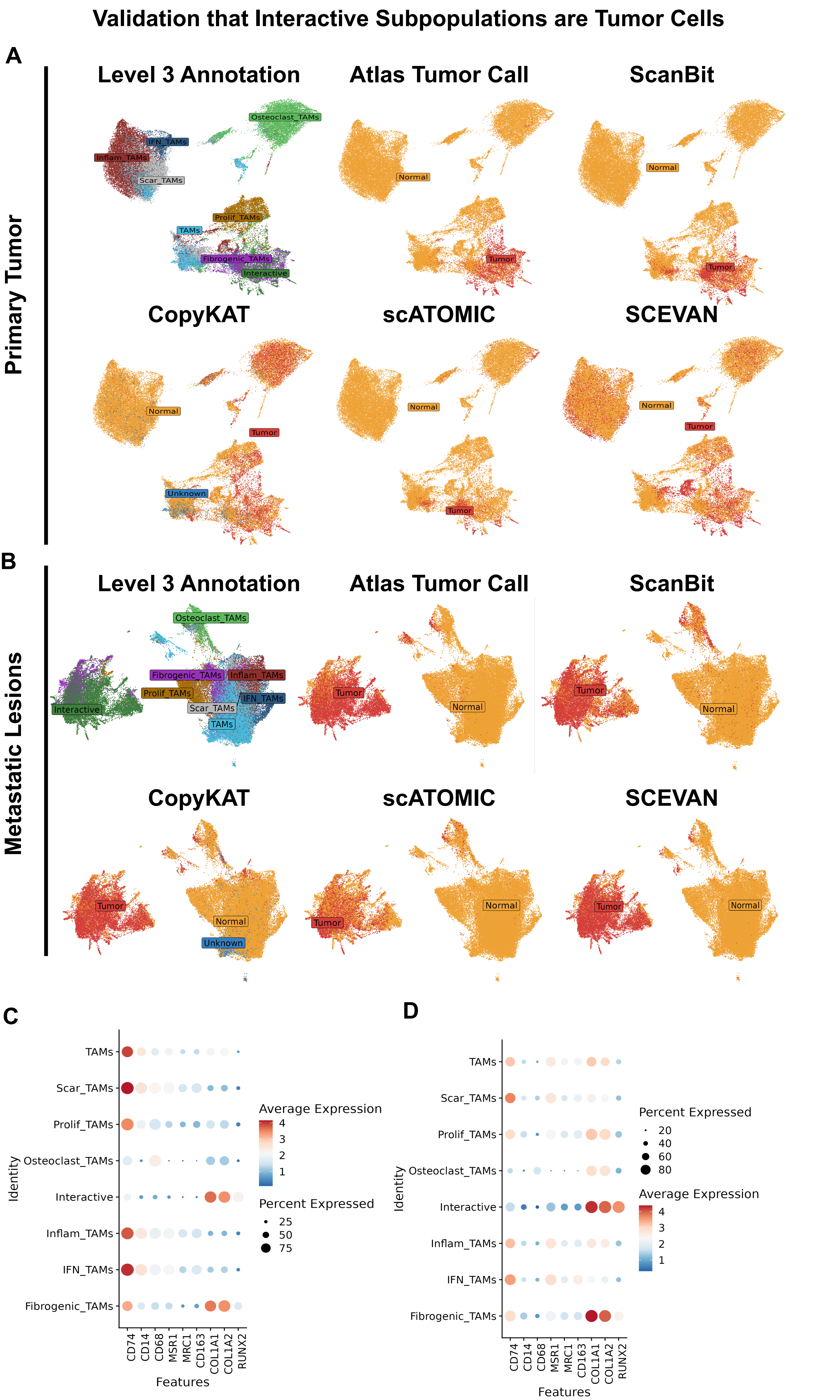


**Supplementary Figure 6 | Analysis of tumor-interactive cells and their relationship with tumor-associated macrophages (TAMs) in human tumors.** Dimensional reduction (DimPlot) visualizations of human patient tumor cells showing A) primary and B) metastatic samples, colored by cell type annotation and tumor–normal classification from the truth-set-based pipeline, scATOMIC, SCEVAN, CopyKAT, and ScanBit. Tumor-interactive cells partially overlap with the fibrogenic TAM subpopulation but form a distinct cluster, indicating a reprogrammed tumor state rather than an immune or TAM phenotype. Dot plots showing selected immune and osteosarcoma tumor markers for C) human patient primary and D) human patient metastatic samples demonstrate that tumor-interactive cells exhibit low expression of immune markers and high expression of tumor markers, supporting their classification as tumor-like rather than immune-like cells.

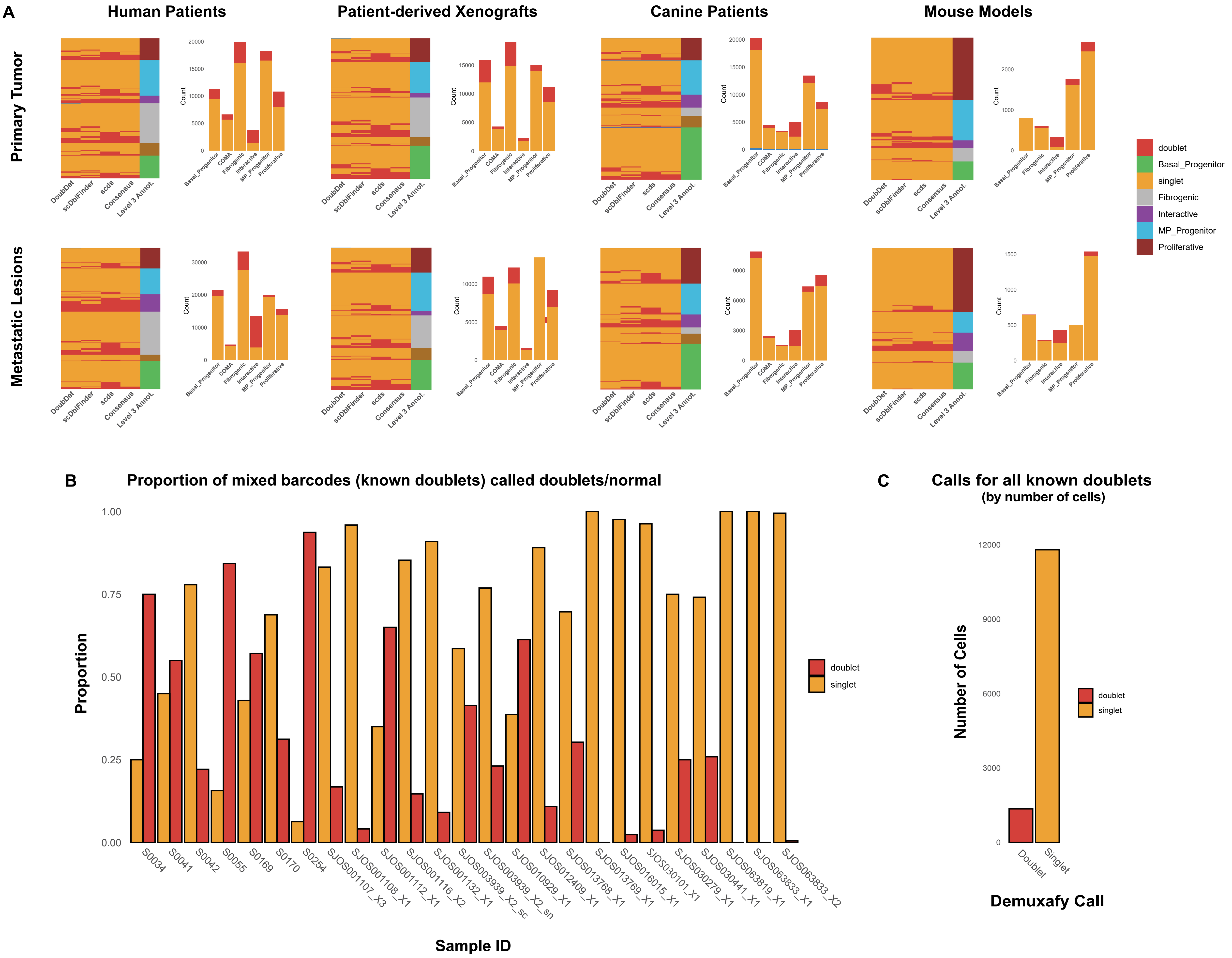


**Supplementary Figure 7 | Doublet identification across tumor subpopulations.** A) Heatmaps depicting tumor cell subpopulations called as doublet or non-doublet across multiple methods employed within the Demuxafy suite. Adjacent barplots show the total number of cells dientified as singlets or doublets by at least two of the three methods for each species and tumor site. In general, the inter-rater consistency of these calls was poor across most cell types. B) Evaluation of doublet detection accuracy using the PDX models to create a truth set. Data show proportion of known doublets (based on balanced human and murine transcript reads) that were called as singlets vs doublets by the algorithm. In general, the sensitivity (and corresponding accuracy) was poor throughout these truth sets. C) Overall performace of the consensus Demuxafy calls in identifying known doublets.

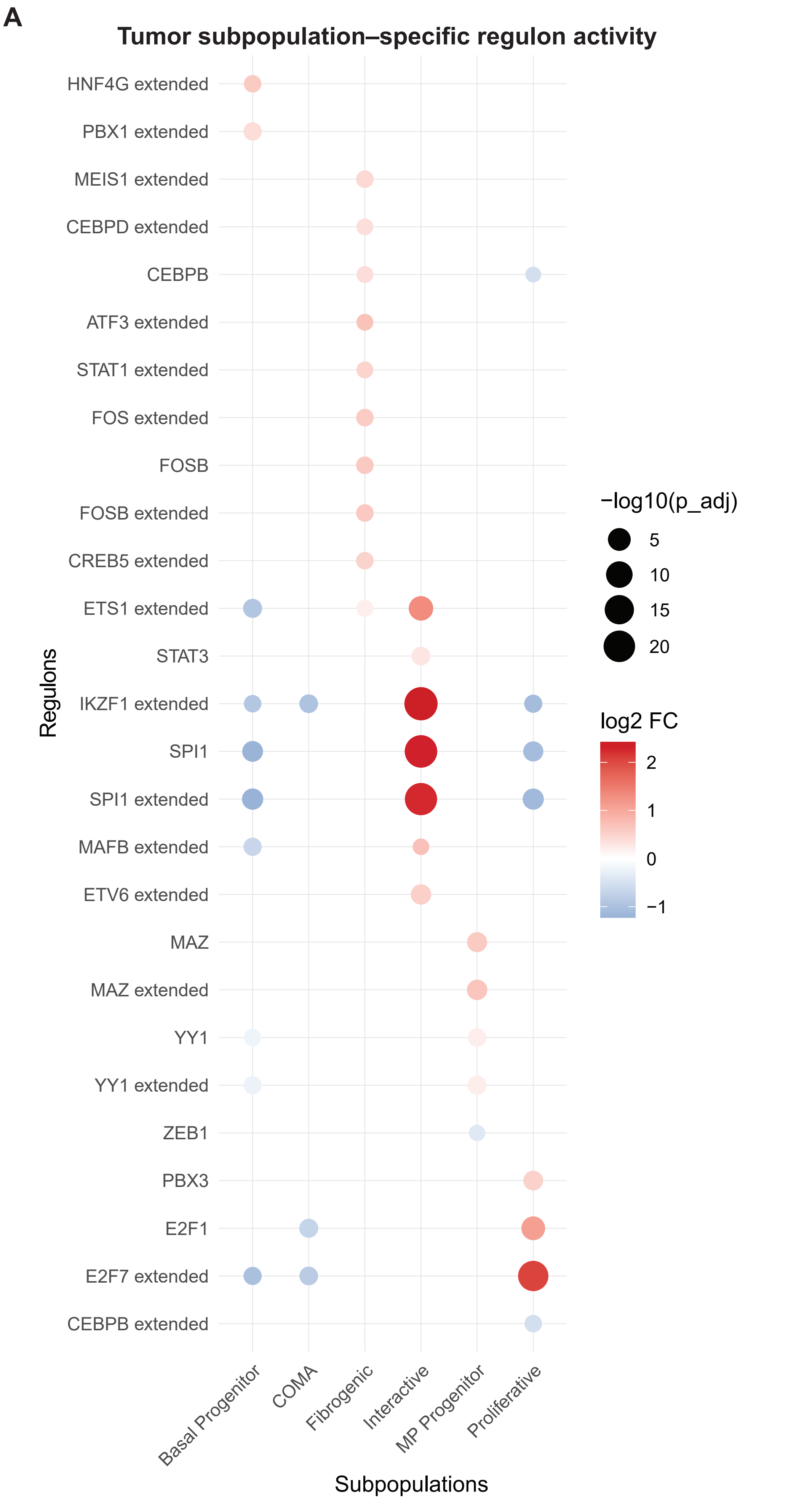


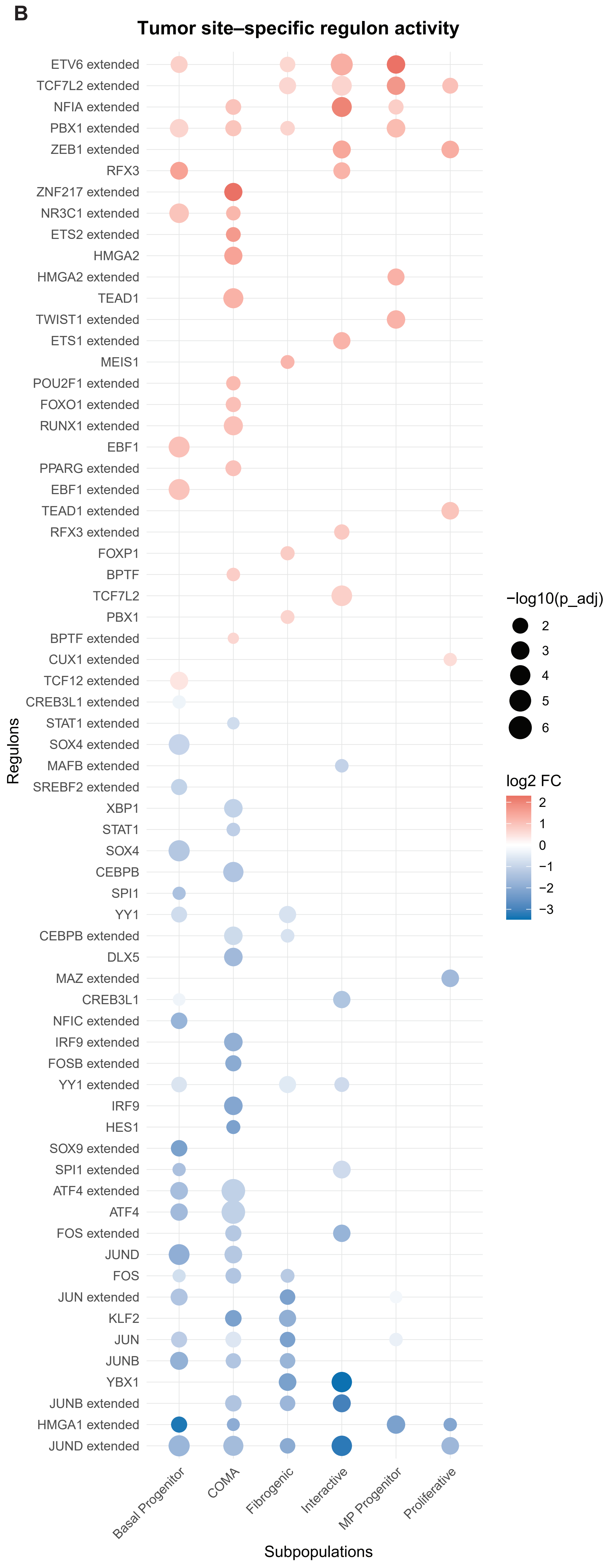


**Supplementary Figure 8 | Regulon analysis using the SCENIC package.** A) Bubble plot showing significantly enriched regulons across tumor subpopulations, independent of tumor site. Regulon activity is compared within each subpopulation relative to all others. Red indicates upregulation and blue indicates downregulation (adjusted $p < 0.05$). B) Bubble plot showing differential regulon activity between primary and metastatic sites within each tumor subpopulation. Red indicates higher activity in metastatic samples, whereas blue indicates higher activity in primary samples (adjusted $p < 0.05$).

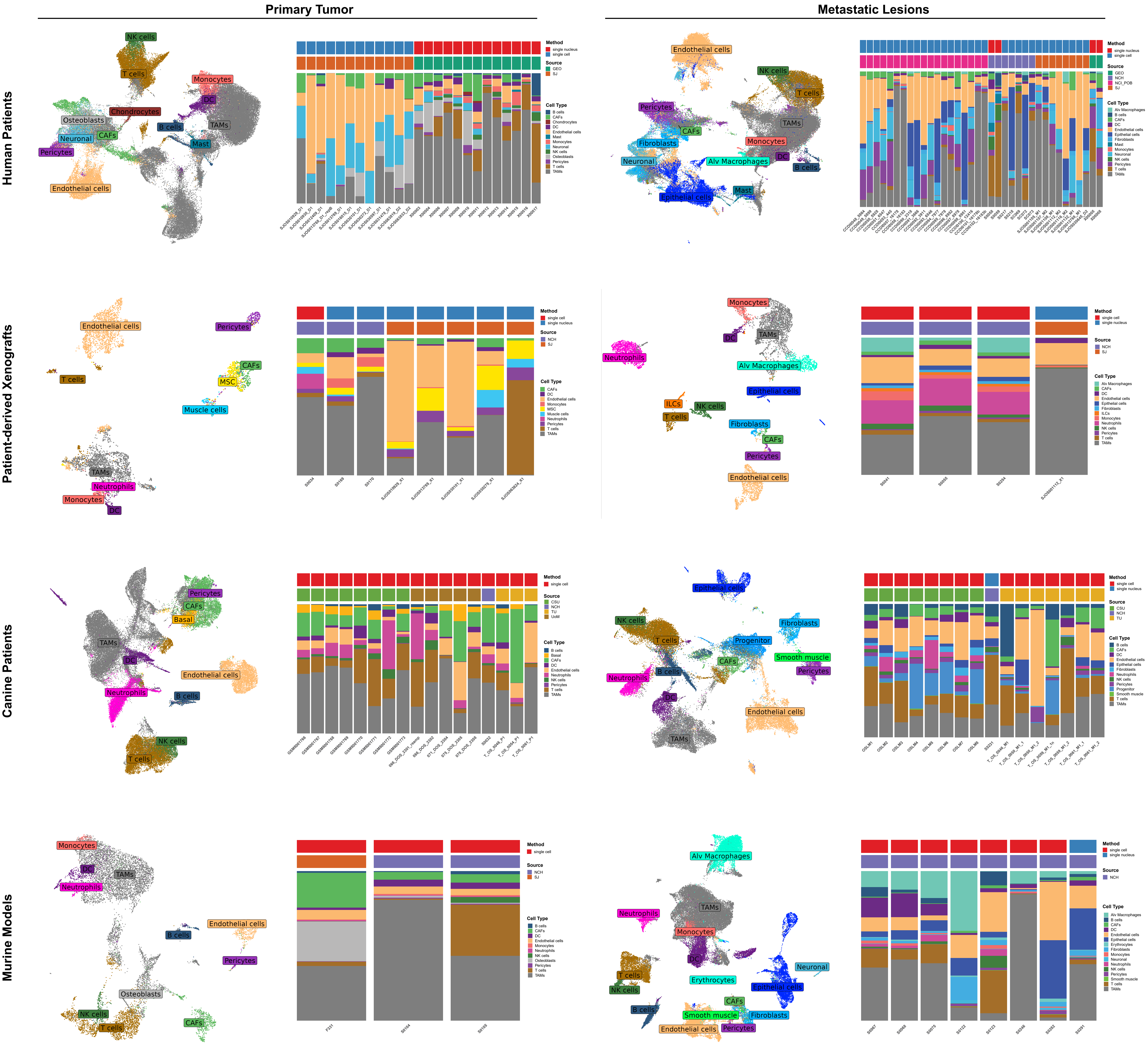


**Supplementary Figure 9 | High-level (Level 2) stromal annotations across sites and species.** DimPlots and stackec barplots showing unified annotations and sample-level composition data for each site and species/model within the atlas. Level 2 annotations correspond to general cell type labels, representing one level below Level 3 annotations, which provide the most granular cell type classification. Integrated lables illustrate the effects of different sequencing methodologies and institutions on the cell types captured within each sample.

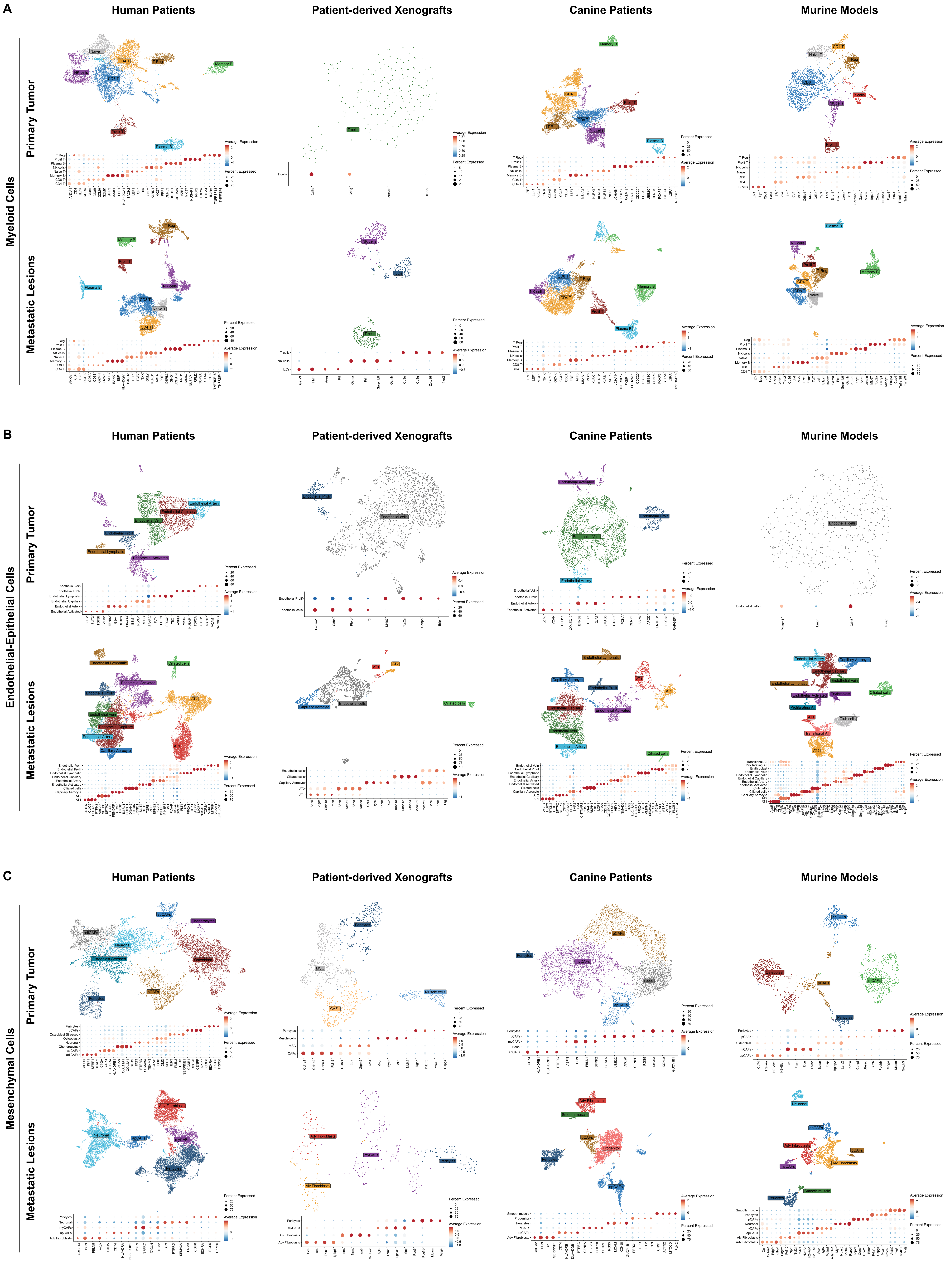


**Supplementary Figure 10 | Detailed subclustering of stromal cell compartments across datasets.** DimPlot and marker dot plot panels depict A) Myeloid, B) Endothelial, and C) Mesenchymal cell compartments across tumor sites and species/models.

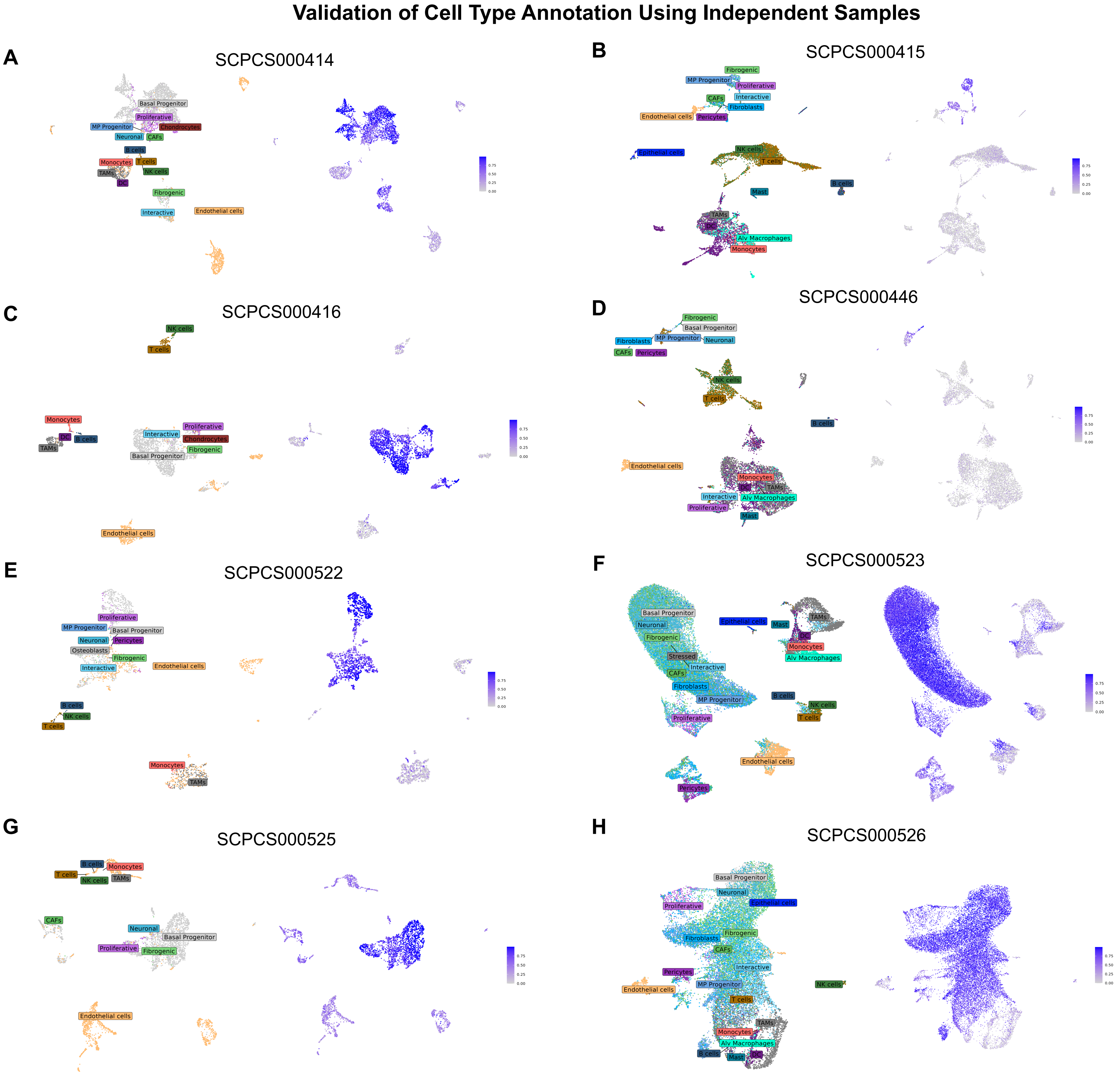


**Supplementary Figure 11 | Validation of the single-cell reference using independent osteosarcoma scRNA-seq datasets.**

Validation of the single-cell reference was performed using publicly available primary osteosarcoma patient samples and metastatic patient-derived organoid datasets obtained from the Alex's Lemonade Stand Foundation repository. Left panels display level 2 cell type annotations transferred from the reference, while right panels show AUCell enrichment scores for canonical osteosarcoma marker genes (RUNX2, SATB2, COL1A1, and COL1A2), demonstrating consistent identification of tumor populations across datasets. A) SCPCS000414, B) SCPCS000415, C) SCPCS000416, D) SCPCS000446, E) SCPCS000522, F) SCPCS000523, G) SCPCS000525, H) SCPCS000526.

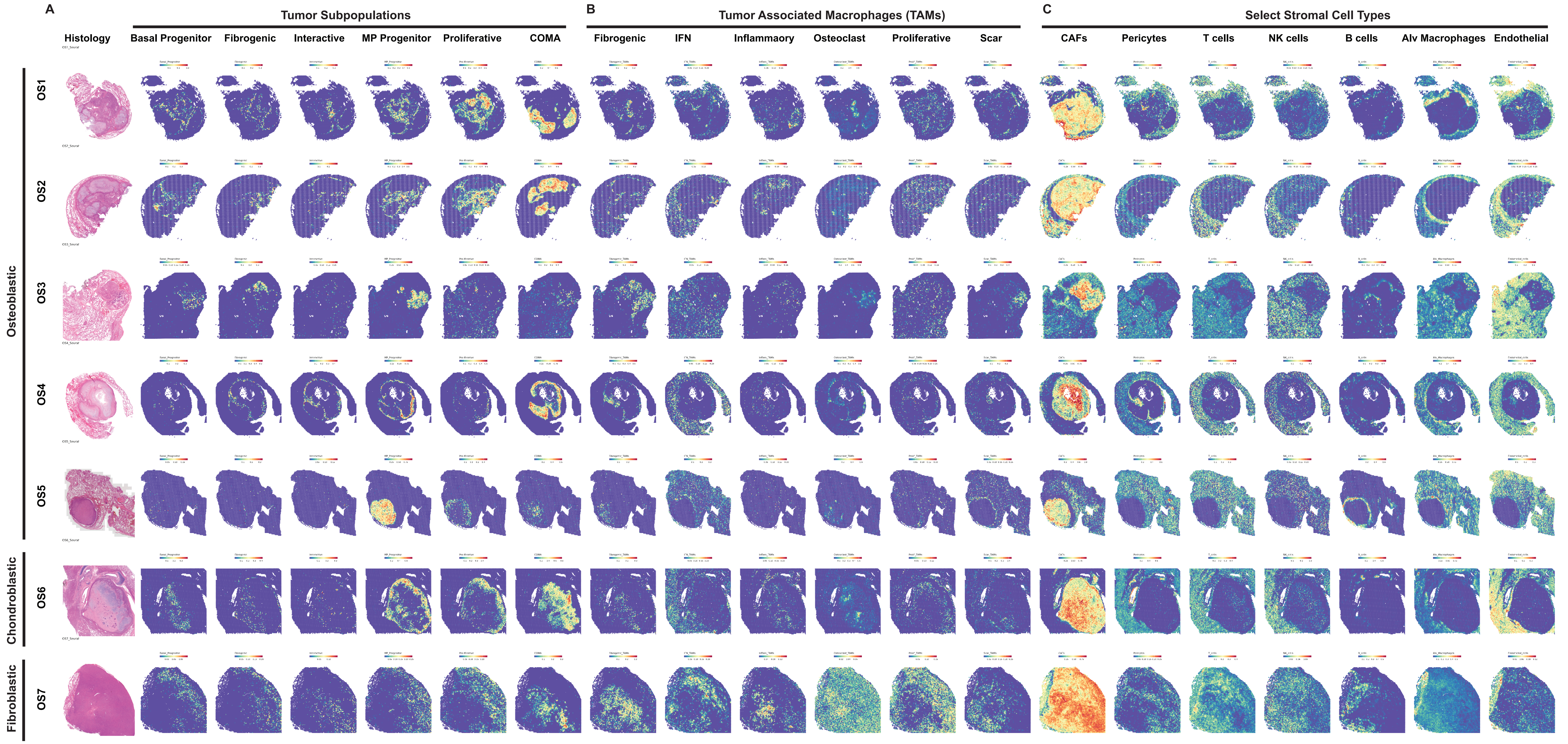


**Supplementary Figure 12 | Spatial deconvolution of cell types in pulmonary samples, highlighting the spatial enrichment and localization of distinct populations.** A) Tumor subpopulations, B) Tumor-associated macrophages (TAMs), C) Select non-tumor cell types.

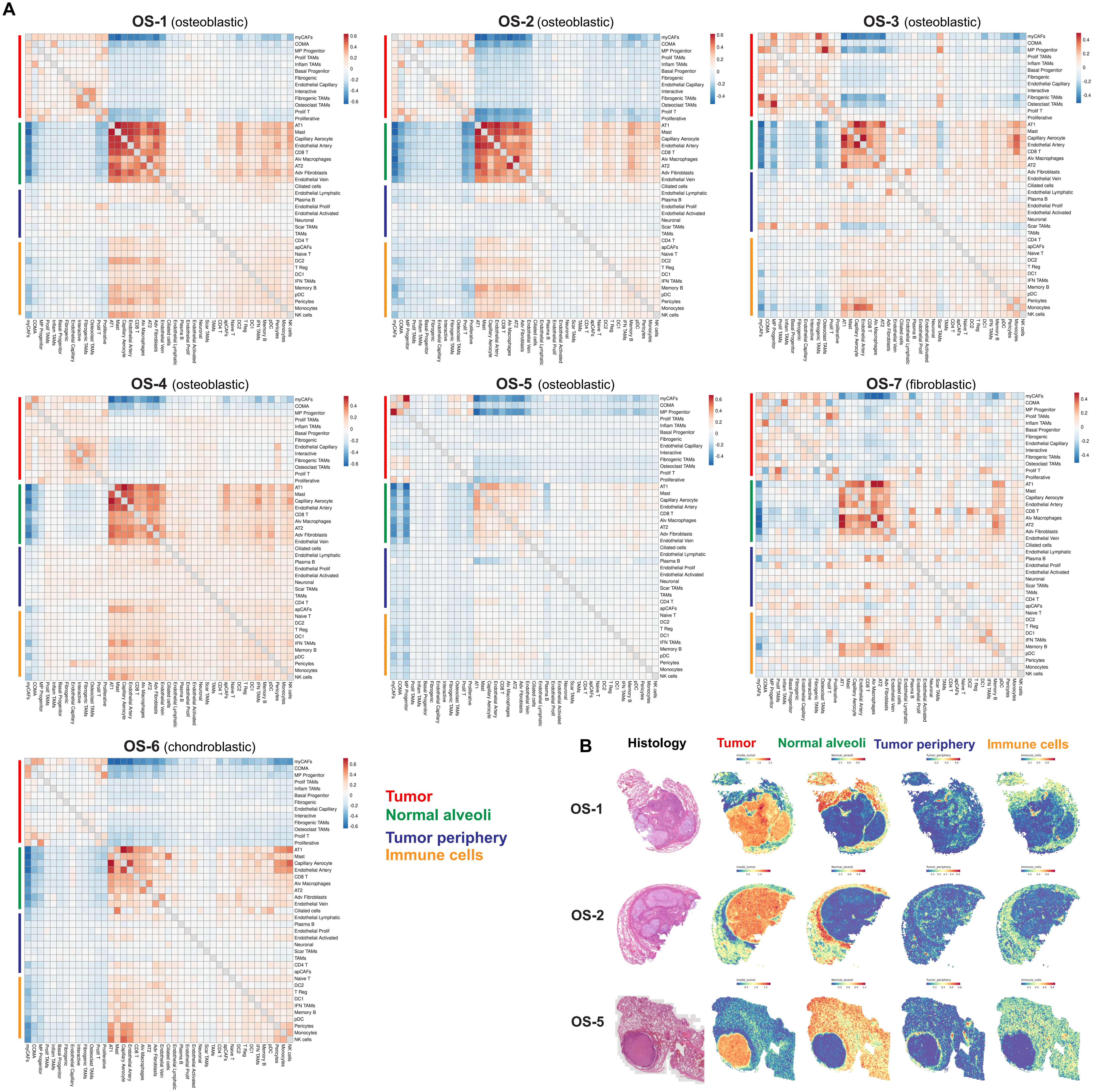


**Supplementary Figure 13 | Neighborhood analysis of cell types in human patient lung samples.** A) Heatmap showing spatial neighborhoods of cell types in each of the Visium dataset. Distinct neighborhoods were identified based on spatial location, including regions within or near the tumor lesion and regions farther away. The Lee statistic was used to quantify spatial co-occurrence relationships between cell types, highlighting replicable patterns of cellular organization within the tumor microenvironment. B) Spatial feature plot of selected samples, showing each neighborhood identified by calculating the summed cell-type features.

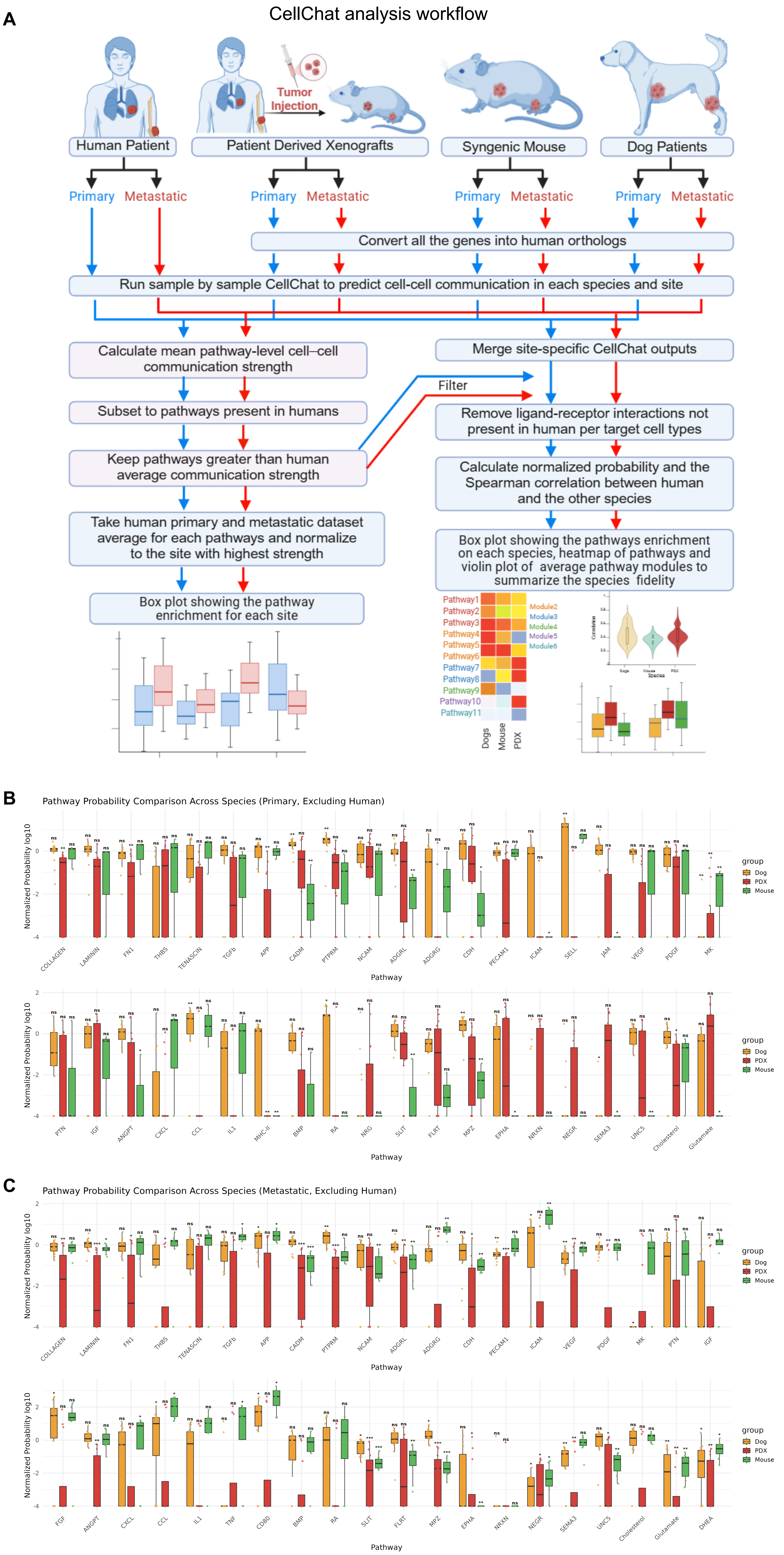


**Supplementary Figure 14 | CellChat analysis workflow and pathway fidelity across species.** A) Schematic illustrating the CellChat analysis workflow and the generation of downstream plots. The diagram outlines how signaling pathways were computed across all datasets by converting genes to their human orthologs, and how significant pathways were subsequently filtered using the human. Boxplots showing CellChat-derived cell–cell communication pathway activity in canine, PDX, and mouse datasets, normalized to the mean pathway activity in human patients for B) primary sites and C) metastatic sites. Each dot represents an individual sample within the respective dataset. P values were calculated by comparison with the human cohort (*P < 0.1; **P < 0.05).
